# Iron Deficiency Impairs Mitochondrial Energetics and Early Axonal Growth and Branching in Developing Hippocampal Neurons

**DOI:** 10.1101/2025.10.15.682603

**Authors:** Daniel C. Mendez, Karishma Devgun, Timothy R. Monko, Luke H. Carlson, Daniel J. Mickelson, Lorene M. Lanier, Michael K. Georgieff, Thomas W. Bastian

## Abstract

Each stage of neuronal development (i.e., proliferation, differentiation, migration, neurite outgrowth and synapse formation) requires functional and highly coordinated metabolic activity to ultimately ensure proper sculpting of complex neural networks. Energy deficits underlie many neurodevelopmental, neuropsychiatric and neurodegenerative diseases implicating mitochondria as a potential therapeutic target. Iron is necessary for neuronal energy output through its direct role in mitochondrial oxidative phosphorylation. Iron deficiency (ID) reduces mitochondrial respiratory and energy capacity in developing hippocampal neurons, causing permanently simplified dendritic arbors and impaired learning and memory. However, the effect of ID on early axonogenesis has not been explored. We used an embryonic mixed-sex primary mouse hippocampal neuron culture model of developmental ID to evaluate mitochondrial respiration and dynamics and effects on axonal morphology. At 7 days in vitro (DIV), ID impaired mitochondrial oxidative phosphorylation capacity and stunted growth of both the primary axon and branches, without affecting branch number. Mitochondrial motility was not altered by ID, suggesting that mitochondrial energy production --- not trafficking --- underlie the axon morphological deficits. These findings provide the first link between iron-dependent neuronal energy production and early axon structural development and emphasize the importance of maintaining sufficient iron during gestation to prevent the negative consequences of ID on brain health across the lifespan.

**Significance Statement:** This study used a primary mouse hippocampal neuron culture model of iron deficiency to address how disruption of iron-regulated mitochondrial activities affects axonal development. After axon initiation but prior to rapid dendrite outgrowth, iron chelation reduced mitochondrial oxidative phosphorylation capacity and stunted the growth of the primary axon and branches but without affecting branch number. Mitochondrial motility was not altered in iron-deficient axons, indicating that reduced neuronal energetic capacity and not impaired axonal mitochondrial trafficking may underlie these morphological deficits. Many neurodevelopmental, neuropsychiatric, and neurodegenerative disorders are characterized by iron and/or mitochondrial dysregulation, highlighting the importance of advancing knowledge on the effects of mitochondrial deficits in early life as it pertains to optimizing brain health throughout the lifespan.

## Introduction

Iron-deficiency (ID) is the most common nutrient deficiency globally and affects 40-50% of pregnant women and children (Gao et al., 2025; Lozoff & Georgieff, 2006). Fetal and neonatal brain development is a period of high iron and energy requirements (Fretham et al., 2011; Georgieff et al., 2018; Radlowski & Johnson, 2013). ID during this early life period acutely impairs brain development and function and increases the long-term risk for cognitive deficits, schizophrenia, ADHD and depression (Insel et al., 2008; Konofal et al., 2004; Lozoff et al., 2013; Lukowski et al., 2010; Schmidt et al., 2014). The hippocampal region of the brain, involved in learning and memory, is rapidly developing during this iron-sensitive period and is particularly iron and energy demanding. Fetal-neonatal ID impairs hippocampal-mediated learning and memory in children, adolescents, and adults even after early postnatal iron repletion (Fretham et al., 2011; Georgieff et al., 2018; Radlowski & Johnson, 2013).

Rodent models of early-life ID have demonstrated reduced hippocampal neuron dendritic arborization and dendritic spine density (Bastian et al., 2016, 2022; Brunette et al., 2010; Carlson et al., 2010; Fretham et al., 2012; Greminger & Mayer-Pröschel, 2015; Jorgenson et al., 2003; Perng et al., 2021), which persist into adulthood even after iron repletion. These dendrite structural deficits on the post-synaptic side of hippocampal neurons may underlie the persistent deficits in synaptic function and learning and memory behaviors following recovery from fetal-neonatal ID. In contrast, very little is known about the effects of fetal-neonatal ID on axonal structural development, outside of hypomyelination (Lozoff & Georgieff, 2006; Todorich et al., 2009). ID has been shown to increase axon diameter and disrupt conduction velocity in the developing auditory nerve, suggesting ID may impair axonal structural maturation (Lee et al., 2012). It is important to advance knowledge on axonal/presynaptic ID effects because axon differentiation and rapid growth/branching of axons occurs much earlier than dendrite growth/branching with major axonal pathways emerging 10 weeks post conception in humans (Žunić Išasegi et al., 2018) and embryonic day 11-18 in rodents during cortical development (Cottam et al., 2025; Dotti et al., 1988). Neurites destined for dendrite specification undergo rapid growth and branch beginning later in development (late gestation - first 2 years after birth in humans and first 2-3 weeks in rodents) (Semple et al., 2013). This also is an important gap in knowledge because during fetal-neonatal brain development, axonal innervation of target regions is often necessary for proper neurogenesis, cell migration and dendrite outgrowth, branching, and spine formation (Andreae & Burrone, 2014; Ghosh & Shatz, 1992; O’Leary & Koester, 1993; Paus, 2023; Yamashita, Aoki, et al., 2016; Yamashita, Yamane, et al., 2016). Previous studies have shown that activity dependent axonal signaling directly influences hippocampal dendritic development through dendritic localization of neurotransmitter receptors, which are critical for activity-dependent dendrite growth and branching (Paus, 2023; Yamashita, Aoki, et al., 2016; Yamashita, Yamane, et al., 2016). If this signaling is disrupted, this results in reduced dendritic branching complexity and synapse formation (Andreae et al., 2014; Paus, 2023; Yamashita, Aoki, et al., 2016; Yamashita, Yamane, et al., 2016). Therefore, misguided axonal growth could contribute significantly to the known structural and functional deficits in the hippocampus following early-life ID.

Iron is critical for cellular energy production through mitochondrial oxidative phosphorylation by providing heme and/or iron-sulfur clusters for cytochrome c and all four protein complexes of the electron transport chain (ETC), as well as succinate dehydrogenase and aconitase in the tricarboxylic acid (TCA) cycle (Bastian et al., 2020). We have previously shown that early-life ID reduces hippocampal neuron mitochondrial respiration and adenosine-5’-triphosphate (ATP) production throughout the period of rapid dendritic arborization (Bastian et al., 2016, 2019, 2020, 2022; de Deungria et al., 2000; Monko et al., 2024; Rao et al., 2003). Neuronal ID also reduces mitochondrial size, motility, and density in the dendritic compartment of hippocampal neurons (Bastian et al., 2019). Thus, we proposed ID-induced dendrite growth and branching deficits are caused by impaired mitochondrial energetics and quality control (Bastian, 2019; Bastian et al., 2016, 2019, 2020). Despite the body of knowledge on mitochondrial and dendritic structure, no studies have addressed the effect of developmental ID on axonal growth and mitochondrial capacity.

Mitochondrial localization and distribution within neuronal sub compartments are critical for neuronal polarization through regulation of ATP production, calcium buffering, reactive oxygen species (ROS) signaling, iron homeostasis and apoptosis (Bradke & Dotti, 1997; Bastian, 2019; Lewis et al., 2013). Inasmuch, the homeostatic balance of mitochondrial dynamics (e.g., bidirectional mitochondria trafficking vs. anchoring) is also important for meeting the metabolic demands of developing axons (Cheng et al., 2022; Cheng & Sheng, 2021; Sheng, 2014). We hypothesized that ID could impair axon development due to iron’s direct role in mitochondrial respiration and the effect of ID on mitochondrial trafficking in dendrites (Bastian et al., 2020). The effects of ID on mitochondrial activity during early neuronal maturation (prior to rapid dendrite outgrowth) remain under-explored.

In the current study, we showed that ID impairs mitochondrial respiration and ATP production rate during the period of rapid axon growth/branching in our hippocampal neuronal culture model of early-life ID. Contrary to our prior dendrite studies, no changes in axon mitochondrial motility were observed in iron-deficient neurons during early axonogenesis. Morphological analysis of iron-deficient axons showed simplified complexity, characterized by stunted growth of both the primary axon and branches. In addition, chronic neuronal ID caused reduced dendritic density of PSD95 puncta. Together, these findings suggest iron-dependent mitochondrial energy production as a key determinant of proper neuronal structural development.

## Materials and Methods

### ANIMALS

Timed pregnant CD1 mice were purchased from Charles River Laboratories. The mice were delivered on embryonic day (E) 13 or 14 and housed in a constant temperature and humidity-controlled facility with food and water administered ad libitum. All animal procedures were performed in facilities accredited by the Association for Assessment and Accreditation of Laboratory Animal Care and in alignment with guidelines and standards established in the *National Institutes of Health’s Guide for the Care and Use of Laboratory Animals.* The protocols for animal procedures were also approved by The University of Minnesota Institutional Animal Care and Use Committee prior to commencement.

### CELL CULTURE

The primary hippocampal neuron culture model of developmental ID used in this study has been previously characterized in long-term studies of mitochondrial function and dendrite growth (Bastian et al., 2016, 2019, 2022; Monko et al., 2024). Briefly, the hippocampi of mixed-sex E16 CD1 mouse embryos were pooled, followed by dissociation and plating at in 6-well plates (125,000 cells/well), 96-well plates (5,000 cells/well), or 24-well XFe microwell plates (8,000 cells/well) coated with poly-D-lysine/laminin. Neuronal plating media [Minimal Essential medium (Corning #15-010-CV), 10mM HEPES (Invitrogen 15630-080), 10mM Sodium Pyruvate (Gibco 11360-070), 0.5mM Glutamine (Invitrogen #25030-164), 12.5µm glutamate (Sigma 128430-100g), 10% fetal bovine serum (Sigma #F6178-500ml), and 0.6% glucose (Fisher MT25037Cl)] was exchanged 2-4 hours after plating cells with glia-conditioned neural growth media [Neurobasal (Gibco #12348-017), 1x B27 (Gibco #17504-044), 0.5mM Glutamine (Invitrogen # 25030-164) and 1x penicillin/streptomycin (Invitrogen #CX30324)]. Time of plating is designated as 0 days *in vitro* (DIV).

At 3 DIV, ID was induced with 5µM of the extracellular iron chelator deferoxamine (DFO, Cayman Chemical #14595) for 24-well XFe (Agilent Technologies) and 96-well plate or 9µM DFO for 35mm dish (Bastian et al., 2016, 2019, 2022; Monko et al., 2024). Concomitantly, 5-fluoro-2’-deoxyuridine (5-FU, Sigma #F0503-100MG) was administered to inhibit glia proliferation during DFO-treatment. Iron-sufficient control cultures received an equal volume of vehicle (ddH2O) in fresh unconditioned neuronal growth medium. As shown previously (Bastian et al., 2016, 2019, 2022; Monko et al., 2024), these DFO doses produce a similar degree of neuronal ID at 11 and 18 DIV as observed in the developing brains of rodent ID models (Carlson et al., 2007, 2009) and human iron-deficient newborn brains (Petry et al., 1992). They do not affect neuronal zinc status (Bastian et al 2016). Assays were performed at 7 DIV since by this time in hippocampal neuronal cell cultures the axonal initial segment (AIS) has been well formed (2-4 DIV) and this point marks the beginning of rapid axonal branching and the initial period of synapse formation, which is critical for proper development of complex neural networks (Grabrucker et al., 2009; Sahu et al., 2019).

### mRNA EXPRESSION ANALYSIS

TRIzol cell lysates from hippocampal cultures at 7 DIV were pooled from 8 wells of a 96-well plate or 1 well of a 6-well plate and RNA was extracted with a Monarch^®^ Spin RNA Cleanup kit (New England Biosciences, catalogue: T2030L). Purity and concentration of samples was determined using a nanodrop spectrophotometer (catalogue #: ND-1000) and 100 ng of RNA diluted in nuclease-free water (20µL reaction volume) was reverse transcribed to synthesize cDNA for quantitative real-time PCR (qPCR) analysis using a cDNA synthesis kit (Applied Biosciences, Catalogue: AB#4387406). TaqMan probe for the *transferrin receptor 1* (*Tfr1*) gene (Thermo, Catalogue: 4331882, Mm00441941_m1) was used as a measure of cellular iron status as we have described (Bastian et al., 2016, 2019, 2022; Monko et al., 2024). TaqMan probes for *TATA binding protein* (*Tbp*) (Thermo, Catalogue: 4331182, Mm00446971_m1), *ribosomal protein s18* (*Rps18*) (Thermo, Catalogue: 4331182, Mm02601777_g1), and *glyceraldehyde-3-phosphate dehydrogenase* (*Gapdh*) (Thermo, Catalogue: 4331182, Mm99999915_g1) genes were used as reference genes for normalization. Briefly, qPCR was performed in triplicate on 2.5ng cDNA using the LUNA master mix (NEB #M3004) adapted from the manufacturer’s protocol for a 10µL reaction volume. An Applied Biosystems QuantStudio 3 qPCR machine (catalogue: A28132) was used with a denaturation step at 95℃ for 1 min., PCR amplification step at 95℃ for 15 sec. and extension at 60℃ for 30 sec per cycle (40 cycles total). Relative gene expression for all genes of interest was calculated relative to an internal calibrator cDNA sample. GeNorm was used to determine the stability of *Tbp*, *Rps18*, and *Gapdh* and their geometric mean was used for normalization of *Tfr1* by ΔΔCt method. Relative *Tfr1* gene expression data from DFO-treated cells was normalized to the average of all IS samples for each unique neuron culture preparation.

### INTRACELLULAR ATP ASSAY

Intracellular ATP concentrations were quantified following manufacturer’s instructions (Promega Cell Glo Titer 2.0) and as reported previously (Bastian et al., 2019, 2022; Monko et al., 2024). Prior to the assay, 1µM Hoechst 33342 was added to each well and live nuclei imaging was performed to normalize ATP concentrations to cell density as we have described (Bastian et al., 2022). Luminescence was measured with a BioTek Synergy LX Multimode Plate Reader. The ATP concentrations of each well were determined based on standard curve readings and then normalized to cell count from live Hoechst imaging (see normalization described below).

### ATP RATE ASSAY FOR BIOENERGETICS

The Seahorse XF ATP Rate Assay was performed according to the manufacturer’s instructions and as described previously (Bastian et al., 2022), to assess relative ATP contribution from either oxidative phosphorylation or glycolysis. In brief, XF Assay Medium (XF DMEM, pH 7.4, with final concentrations of 1mM XF Sodium Pyruvate, 2mM XF Glutamine, and 10mM XF Glucose) was prepared and sterile filtered either the day before and stored at 4°C or prepared the same day. On the day of assay at 7 DIV, neuronal medium was replaced with XF Assay Medium containing either 5µm DFO or vehicle. Working dilutions of compounds, Oligomycin (ATP synthase inhibitor) and Antimycin A/Rotenone (Complex III and I inhibitors, respectively) were made to achieve a final concentration of 1.5µM and 0.5µM after injection, respectively, in XF Assay Medium. To de-gas the plate of CO_2_, the cell culture microwell plate was placed immediately in a 37°C non-CO_2_ incubator for 30 minutes (min) prior to the start of the bioenergetics assay. Oxygen consumption rates (OCR) and extracellular acidification rates (ECAR) were measured simultaneously and normalized to cell density as done previously and described below (Bastian et al., 2022).

### INTRACELLULAR AND ATP RATE CELL DENSITY NORMALIZATION

To normalize intracellular ATP and Seahorse OCR and ECAR measurements to cell density, neuron cultures were treated with 1uM Hoescht 33342 (Cayman Chemical, 15547) for live nuclear imaging and placed in humidity and temperature-controlled incubation (5% CO2 and 37°C) for live cell imaging using Zeiss Celldiscoverer 7 and Zeiss Axiocam 506 mono camera with the following specifications: 5x Plan-Apochromat objective (NA=0.35), LED illumination for oblique contrast transmitted light (725nm) and fluorescence (385nm), and fluorescent quad bandpass filter set 90HE (beamsplitter: RQFT 405 + 493 + 575 + 653nm, emission: QPB: 425/30 + 514/30 + 592/25 + 709/100nm). Data for intracellular and ATP rate assays were normalized to the number of cells in the respective well. Images were segmented, classified, measured, and mapped to experimental data in batch using tools from our Python package, [napari-ndev] (T. Monko & Bastian, 2025). Images were loaded into Python using [bioio](https://github.com/bioio-devs/bioio) and [pyclesperanto] (Haase et al., 2023). GPU-accelerated algorithms were used to pre-process by first reducing noise (median) and then subtracting background (tophat). Images were segmented with pyclesperanto by using [Otsu’s] (Otsu, 1979), thresholding algorithm on gaussian blurred images followed by separation of neighboring cells with a seeded watershed, created by a voronoi diagram of peak spot detection of a gaussian blurred image. To detect live and dead cells among the segmented nuclei, we trained Random Forest machine learning object classifiers using napari-ndev and [accelerated pixel and object classifiers] (Haase et al., 2022). Segmented nuclei were predicted for nuclei classification in batch with napari-ndev. Finally, segmented images were measured with napari-ndev tools using either [scikitimage] (Van Der Walt et al., 2014) or [Nyxus] (https://github.com/PolusAI/nyxus) as the label measurement backend. Image metadata, including well position in the respective culture plate, was used to map data to treatment conditions and other experimental metadata using napari-ndev’s PlateMapper. Data was grouped and summarized according to this metadata. Intracellular ATP data values were relativized to the average control value for each unique culture experiment and log transformed before statistical analysis.

### MITOCHONDRIAL MOTILITY LIVE IMAGING

Prior to plating, cells were co-nucleofected with an mcherry-Mito-7 (Addgene # 55102) and “humanized” recombinant green fluorescent protein (hrGFP) plasmid driven by a CMV enhanced chicken B-actin promoter (Zolotukhin et al., 1996), using a Lonza Amaxa II and Mouse Neuron Nucleofector Kit (Cat. No. VAPG-100), as described previously (Bastian et al., 2016; Penrod et al., 2015). Nucleofected cells were mixed 1:1 with non-nucleofected cells and plated in 96-well plates at a seeding density of 5,000 cells/well. At 7 DIV, time-lapse imaging was performed using a ZEISS Celldiscoverer 7 with environmental control set to 37^°^C and 5% CO_2_. mCherry-positive neurons (n = 11-12 neurons per group) were imaged in 3-channels (oblique, mcherry and hrGFP) every 1 second (s) for a duration of 1.7 min for one independent time-lapse experiment or 3.5 s for 5 min for a second independent time-lapse experiment with a ZEISS Axiocam 506 mono camera with the following specifications: 50x Plan-Apochromat objective (NA=1.2) with water immersion and 0.5x magnification, LED illumination for oblique contrast transmitted light (725nm), and fluorescence (590nm & 470nm), and fluorescent quad bandpass filter set 90HE or triple bandpass filter set 91HE (beamsplitter RTFT 450 + 538 + 610; emission filter TBP 467/24 + 555/25 + 687/145).

The images were converted to TIFF files and then organized according to the hierarchical organization described in the KymoAnalyzer manual (Neumann et al., 2017). Axonal segments between 50-100µm were traced anterogradely to the axonal growth cone. Kymographs for each neuron generated were then traced and processed through semi-automated Fiji plugins of the KymoAnalyzer plugin package version 1.01 (Neumann et al., 2017; Schindelin et al., 2012). Data for the various mitochondrial parameters were then organized by category, mean and median compiled for individual mitochondria for each imaged axon, then analyzed and graphed in PRISM.

### IMMUNOCYTOCHEMISTRY OF AXONAL PROCESSES

Cells nucleofected with hrGFP were mixed 1:1 with non-nucleofected cells and plated in a 6-well plate (125,000 cells/well). This allowed sparse neuronal labeling to trace individual neurons. At 7 DIV, cultures were fixed and processed for immunocytochemistry (ICC) in 6-well plates as described (Bastian 2016, 2017, 2019, 2022). Briefly, cultures were incubated with a mouse monoclonal primary antibody against MAP2 (Abcam, Catalogue: ab11268) at a 1:500 dilution for 1 hour at room temperature (RT) or overnight at 4℃ to specifically label the dendrites. Cells were then incubated with a secondary donkey anti-mouse Alexa Fluor^®^ (AF) 647 antibody (Jackson Immunoresearch, Catalogue:715-605-151) diluted at 1:50 for 1 hour at RT. DAPI stain was used to visualize the nucleus. Individual GFP-positive neurons were imaged using a ZEISS Celldiscoverer 7 and Zeiss Axiocam 506 mono camera with 20x Plan-Apochromat objective (NA=0.7) at 0.5x magnification and fluorescent filter sets described above. Tiled images were stitched using ZEISS Zen Software and exported as OME-TIFF for morphology analysis.

### AXONAL MORPHOLOGY ANALYSIS

The images were analyzed using Fiji and the Simple Neurite Tracer plugin to determine the structural effect of iron deficiency on axon length and branching as done previously for dendrites (Bastian et al., 2016, 2022). The center of the soma was identified and marked. In addition to being much longer, thinner and more highly branched than dendrites at this stage, the axon was identified as being GFP-positive and MAP2-negative. The primary axon was determined as the longest and straightest axonal process, with secondary branches emanating directly from the primary axon and all other branches considered tertiary axon branches. Sholl analysis was performed to determine the number of crossing points as a function of distance from soma (overall complexity), and the number of branches and length of primary axon and branches were quantified.

### POST-SYNAPTIC DENSITY ANALYSIS

To enable visualization of endogenous PSD95 in individual neurons, a plasmid expressing an eGFP-tagged intrabody specific for PSD95 (pCAG_PSD95.FingR-eGFP-CCR5TC Addgene plasmid # 46295; a gift from Don Arnold (Gross et al., 2013) was nucleofected into hippocampal cells as previously described (Bastian et al., 2022). Nucleofected cells were then mixed 1:1 with non-electroporated cells before plating on 12mm coverslips in PDL/laminin-coated 35mm petri dishes (150,000 cells per dish) and culturing as described above and previously (Bastian et al., 2016, 2017, 2019, 2022). At 18DIV, hippocampal cultures were fixed in 3.7% paraformaldehyde, and coverslips were mounted on slides as described (Bastian et al., 2016, 2017, 2019, 2022). Terminal dendrite segments of individual eGFP-positive neurons were imaged with a ZEISS Axiovert 200M microscope with 100x plan-neofluar objective (NA=1.3), ZEISS Axiocam HRm CCD camera, Chroma eGFP/FITC filter cube (#49002), and MicroManager software. Z-stack images were taken from the top to the bottom focal plane of the dendrite at a 0.35µm imaging interval and PSD95 puncta density (as a surrogate for spine/synapse density) was quantified using Fiji (Schindelin et al., 2012). A custom ImageJ macro (see supplementary material for code) was developed to automate batch analysis of all neurons. Briefly, z-stacks were merged using a max intensity projection, processed using a custom convolution kernel, and thresholded with the “Mean dark” AutoThreshold feature. Thresholded puncta were counted using “Analyze Particles” and density was calculated relative to dendrite length. Dendrite segment lengths were determined manually using the “Segmented Line” and “Measure” features in Fiji.

### STATISTICAL ANALYSIS APPROACH

Graphpad Prism 10 was used to carry out univariate statistical analyses with α=0.05 to determine statistically significant differences between groups. Unpaired t-test or Mann-Whitney was used for parametric or non-parametric data, respectively for statistical analysis of ATP concentrations, Seahorse respiratory parameters, mitochondrial motility, and PSD95 puncta density. All datasets were tested for normality (Anderson-Darling, D’Agostino-Pearson omnibus, Shapiro-Wilk, & Kolmogorov-Smirnov) and equal variance (F-test) prior to subjecting to parametric or non-parametric tests. If data was normally distributed but had unequal variance, an unpaired t-test with Welch’s correction was applied. To adjust for multiple comparisons, the Benjamini-Hochberg method of FDR correction was used on resulting p-values from Mann-Whitney tests for all mitochondrial motility parameters. Statistics for Sholl Analysis were computed using repeated-measures 2-factor ANOVA to compare group differences with respect to distance from the soma. Multivariate analysis was coded using Python Jupyter Lab Notebook using exported data from all Sholl Analysis parameters. Prior to analysis, Levene’s test showed equal variance but non-normal distribution of the Sholl dataset with or without log-transformation. A Box’s M test also showed differences in covariance matrices between IS and ID groups. Though assumption violations were present due to non-normal distribution and difference in covariance matrices, a multivariate analysis of variance (MANOVA) was still performed which included statistical tests by Pillai’s trace and Wilk’s lambda. Pillai’s trace is a MANOVA statistical approach robust to violations of normality and covariance particularly for datasets with large and balanced sample sizes, while Wilk’s lambda is more sensitive to these assumption violations and outliers (Appolus EE & Okoli CN, 2022). Thus, both statistical results are reported.

## Results

### Neuronal reduction of total intracellular ATP after iron chelation during early axonogenesis

To evaluate 7 DIV neuronal iron status in our hippocampal neuron culture model - as done previously at 11, 18, and 21 DIV (Bastian et al., 2016, 2022; Monko et al., 2024) - mRNA levels of the *transferrin receptor 1* gene (*Tfr1*), which encodes the surface receptor chiefly responsible for cellular iron uptake and is a sensitive indicator of neuronal iron status, were measured by qPCR (Carlson et al., 2007, 2009). Relative mRNA expression of *Tfr1* significantly increased in ID hippocampal neuronal cultures by 1.83-fold in a 6-well vessel using a 9µM DFO dosage and 1.32-fold increase in the 96-well format using a 5µM DFO dosage (Figure 1B). Total intracellular ATP (p = 0.0276) was significantly decreased by 18% in 5uM DFO-treated neurons compared to IS (Figure 1C), consistent with previous work demonstrating reduced energy status at later stages of neuronal development (Bastian et al., 2019, 2022; Monko et al., 2024). ATP measurements were normalized to cell density which was not significantly different between IS and ID wells (p=0.894).

**Figure 1:**
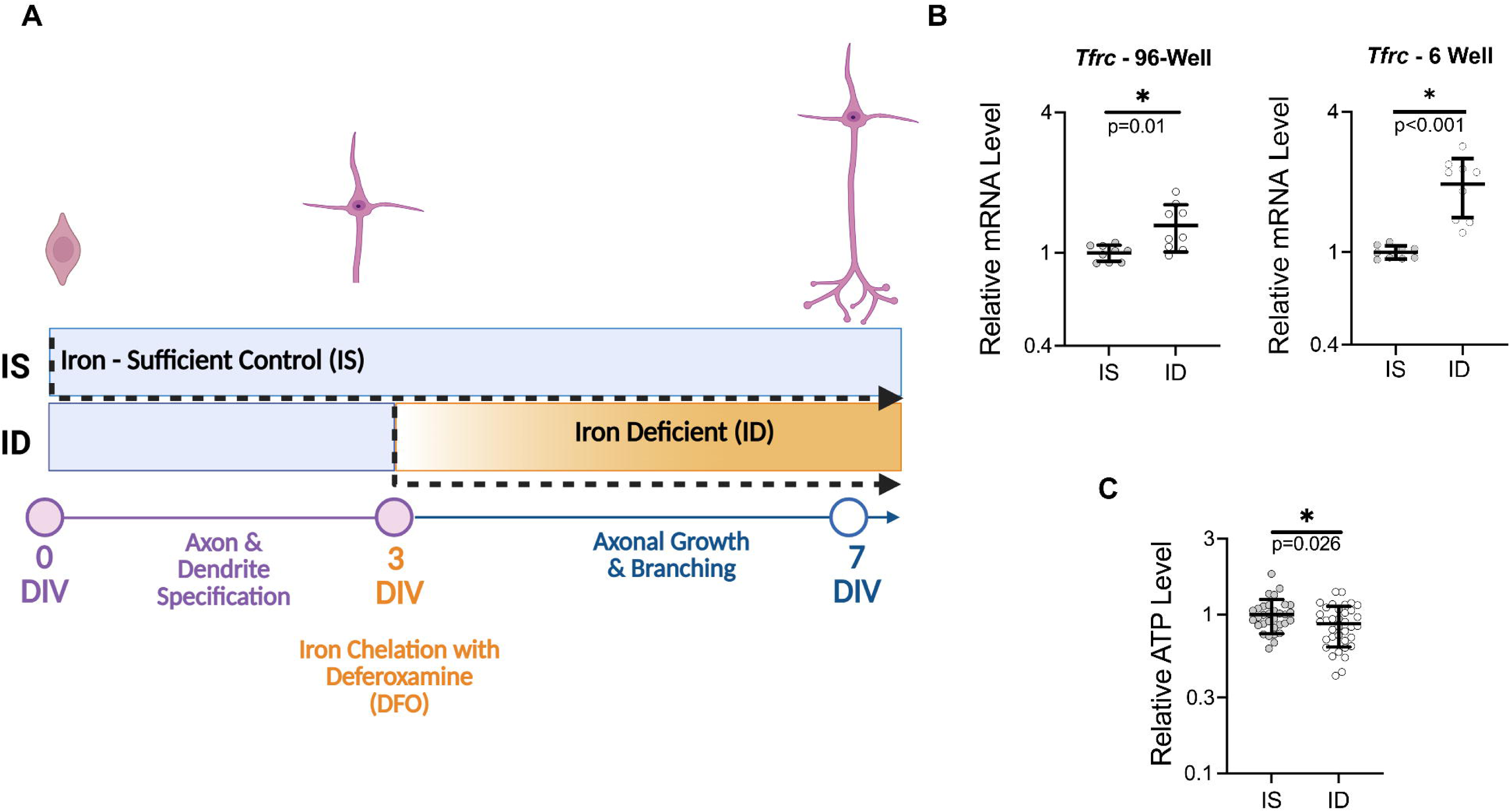
ID decreases total intracellular ATP concentrations in 7 DIV hippocampal neurons during axonogenesis. **A)** Illustration of hippocampal neuron culture experimental design (Created in BioRender). Primary hippocampal neuronal cultures from E16 mice were treated with 5-FU and with or without DFO at 3 DIV. **B)** At 7DIV, quantitative real-time PCR (qPCR) was performed for *Tfrc*, the gene coding for the main neuronal iron uptake protein, to validate iron status of treatment groups (n= 6 independent wells per group pooled from two unique culture preparations). *Tfr1* mRNA levels were calculated relative to the geometric mean of three reference genes (*Tbp*, *Rps18*, and *Gapdh*) and relative change in *Tfr1* mRNA is compared from experiments in 6-well and 96-well vessels. **C)** Total intracellular ATP concentration was measured and normalized to cell count per well (n = 32-36 independent wells per group). Statistical significance was determined on log10-scaled data by unpaired t-test with or without Welch’s correction for qPCR in 96-well and 6-well, respectively, comparing ID induced DFO dose to IS neurons and displayed on linear scale. The parameteric unpaired t-test was used on relativized and on log10-scaled intracellular ATP assay for determining significance between IS versus ID neurons. Statistical significance is denoted with asterisk (*) for p-values ≤ 0.05. For both B and C individual data points representing one independent replicate are calculated relative to the average for the IS group and are shown along with the mean ± SD. Relative data are shown on a log scale to accurately reflect the magnitude of changes.

### Effects of iron chelation on neuronal energetics during axonogenesis

Mitochondrial respiration by oxidative phosphorylation was impaired in 7 DIV ID neurons across several energetic respiration states compared to IS neurons. Specifically, OCR associated with basal respiration (p = 0.011; Figure 2B), ATP-linked respiration (p = 0.025; Figure 2C) and non-mitochondrial respiration (p=0.012; Figure 2D) were decreased by 27%, 25%, and 47%, respectively. This was associated with trending decreases in mitochondrial ATP production rate (p = 0.093) and glycolytic ATP production rate (p=0.108) and a 26% lower Total ATP production rate (p=0.015) in ID compared to IS neurons (Figure 2E-G). ATP rate index (p=0.609) was not different between IS and ID neurons (Figure 2H), which is consistent with previous observations (Bastian et al., 2022). All bioenergetic read-out measurements were normalized to cell density, which was not significantly different between groups (p=0.637).

**Figure 2:**
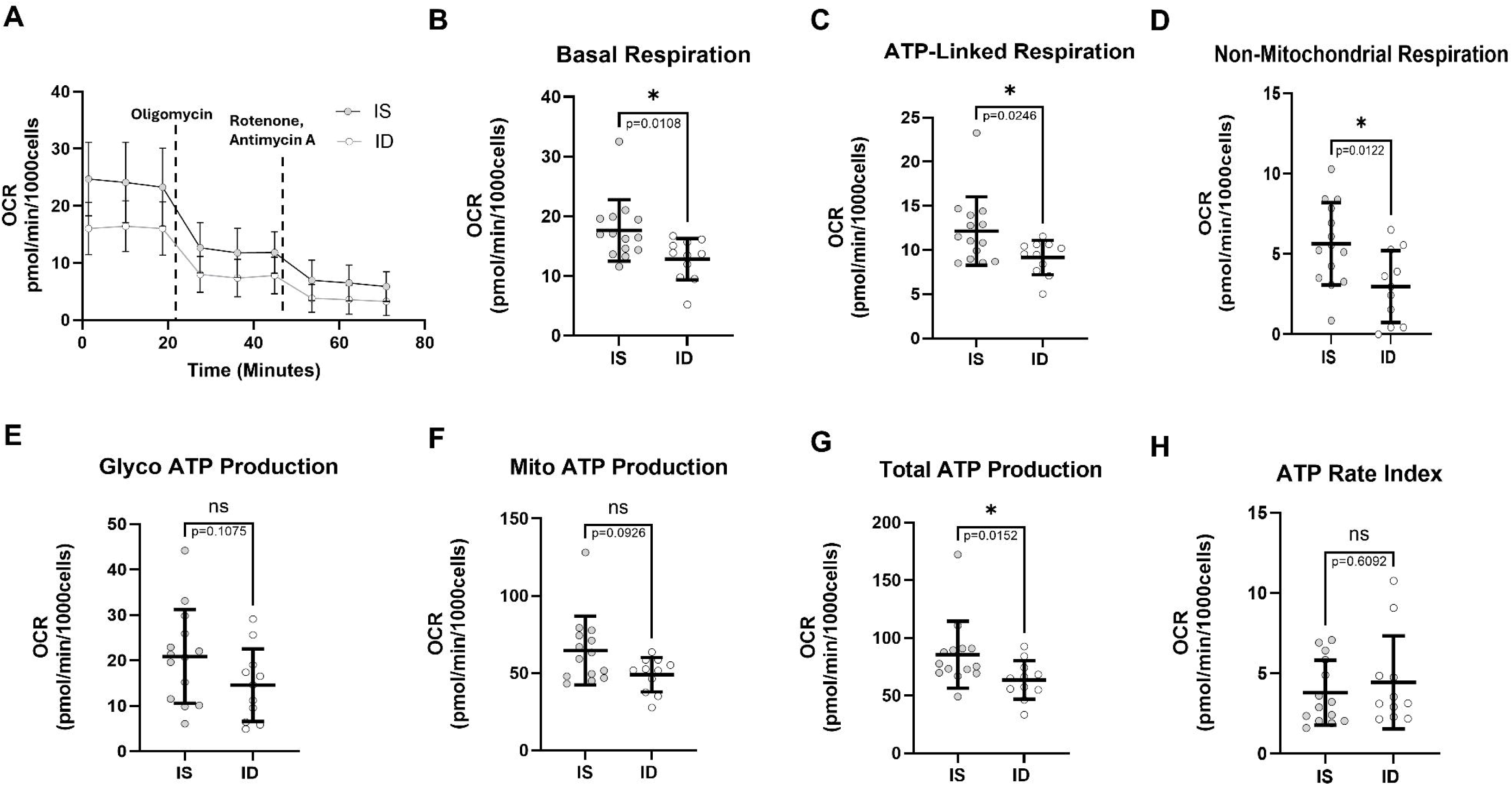
ID impairs mitochondrial respiratory capacity in 7 DIV hippocampal neurons. Primary hippocampal neuronal cultures from E16 mice were treated with 5-FU and with or without DFO at 3 DIV after axon specification and assessed for their respiratory capacity using ATP Rate Assay. **A)** Kinetic plot of real-time OCR normalized to cell count (pmol/min/1000 cells) measured before and following injections of Oligomycin and Antimycin A/Rotenone (AA/Rot). **B-D)** Normalized respiration parameters for non-mitochondrial respiration, basal respiration, and ATP-linked respiration were derived from the real-time OCR data. **E-F)** OCR and ECAR data was used to calculate the ATP production rate from either mitochondrial oxidative phosphorylation or glycolysis. **G)** The sum of both ATP production rates was used to calculate the total ATP production rate. **H)** The ATP Rate Index was calculated by the ratio of mitochondrial ATP production (mito ATP) rate over the glycolytic ATP (glyco ATP) production rate. Data were pooled from three unique culture preparations (n=11-14 independent wells per group). Data points for values of independent wells with the mean ± SD are displayed on each plot. An asterisk (*) indicates a statistically significant difference (p≤0.05) by parametric unpaired-test or non-parametric unpaired Mann-Whitney test for each respiration parameter.

### Iron chelation does not affect mitochondrial motility during axonogenesis

Since mitochondrial trafficking to sites of high energy demand is required to metabolically support developing neuronal processes and is itself an ATP-consuming process, and because we saw mitochondrial motility differences in dendritic compartment of hippocampal neurons due to ID (Bastian et al., 2019), mitochondrial motility was assessed in the axonal compartment (Millecamps & Julien, 2013). Kymograph analysis on time-lapse images (Movie 1A-B) of iron-sufficient and iron-deficient axons (Figure 3 A-B) showed no significant difference in any of the motility parameters (Table 1) including net velocity, segmental velocity (segments of mitochondria motion excluding pauses), density, total pause frequency in retrograde and anterograde direction and percent time in motion (Figure 4A-D, Table 1). However, assessment of sub-categorized data showed a 38.2% decrease in segmental velocity in the retrograde direction in iron-deficient compared to iron-sufficient neurons (p = 0.031; Figure 4B). Specifically, mitochondria moving in the anterograde direction towards the axon terminal paused 57% more in iron-deficient neurons than iron-sufficient neurons (p = 0.006) (Figure 4D). Ultimately, when p-values were adjusted for false discovery rate (FDR) using the Benjamini-Hochberg method, no significant difference was observed for any of the parameters assessed. Similar trends and no significant differences were found for axonal mitochondrial motility parameters when assessed in the 9µm DFO dose/35mm dish culture system (data not shown).

**Figure 3:**
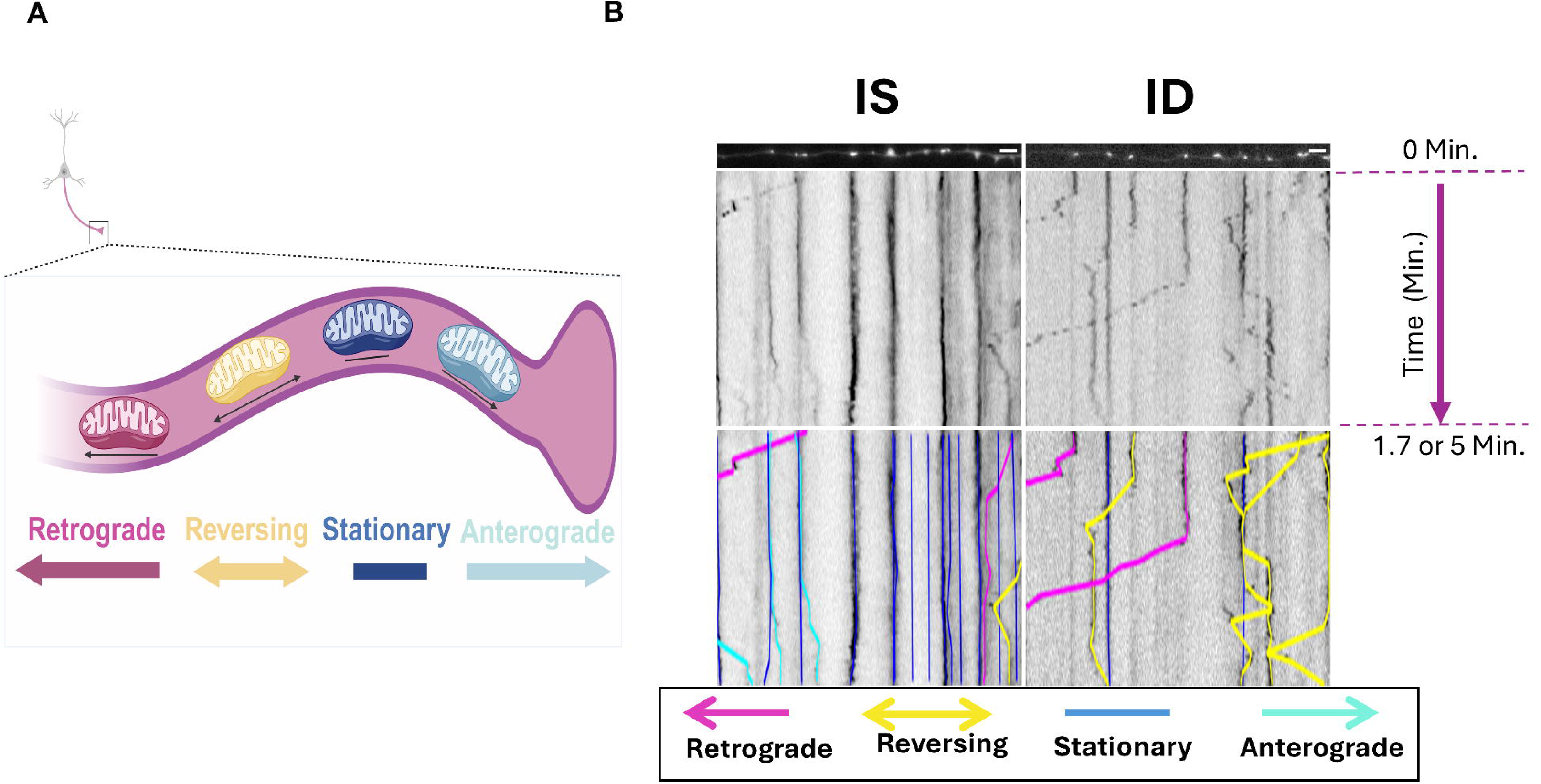
Axonal mitochondrial motility kymograph analysis in 7DIV hippocampal neurons. Primary hippocampal neuronal cultures from E16 mice were treated with 5-FU and with or without DFO at 3 DIV. At 7 DIV, time-lapse imaging (1s or 3.5s intervals for 1.7 or 5min) was performed to capture motility of fluorescently labelled mitochondria in axonal compartment of neurons transfected with mcherry-Mito-7. **A)** Illustration of axonal region of interest analyzed and mitochondrial trafficking activity (created in BioRender). B) Representative still frame (t=0 min) of fluorescently labelled axonal mitochondria is shown with a representative kymograph of IS and ID. To better visualize the traced axonal path, the images were straightened using ImageJ “Straightened” tool. Tracks in kymographs represent individual mitochondria over time (x-axis) and direction (y-axis). Manually traced tracks are color-coded by category of direction of mitochondrial motion relative to the soma and are defined as retrograde (magenta), reversing (yellow), anterograde (cyan), and stationary (blue). Scale bars represent 5µm.

**Figure 4:**
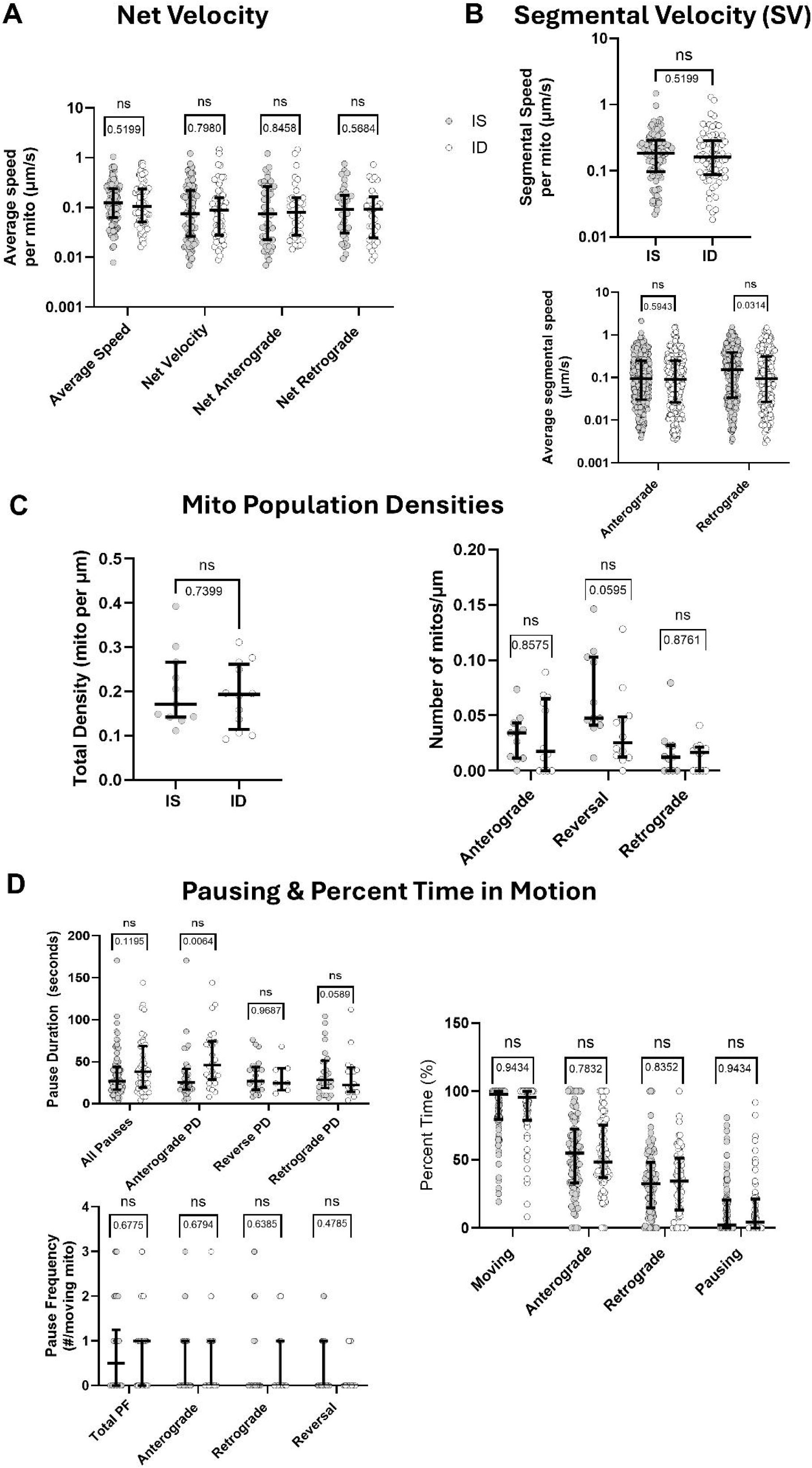
ID does not alter mitochondrial motility in 7DIV axons. **A)** Average speed for all moving mitochondria (regardless of direction or net displacement) was calculated. The net velocity represents any mitochondria (IS: n=98; ID: n= 70) displaying net movement in the anterograde or retrograde direction and displayed as a scalar value (speed µm/s). **B)** The average segmental speed was calculated for each mitochondria only when in active motion (excluding pauses) (IS: n=98; ID: n=70). All individual moving segments were pooled and sub-categorized by anterograde (IS: n=486; ID: n=364) and retrograde (IS: n=391; ID: n=267) directionalities to calculate average segmental speed. All values are represented in scalar form (µm/s). **C)** The total density of mitochondria per axonal length and the density of sub-populations of mitochondria that showed overall anterograde, reversal, or retrograde movement were plotted for IS and ID axons (IS: n=11; ID: n=12). **D)** The pause frequency (PF) of all pausing instances (IS: n=98; ID: n=70) of individual mitochondria for IS and ID was plotted. The PF of individual mitochondria with overall anterograde (IS: n=84; ID: n=58), reversing (IS: n=58; ID: n=31), or retrograde (IS: n=72; ID: n=43) directionality was also extracted and plotted. Time duration in seconds (s) for all pauses (IS: n=79; ID: n=49) was calculated and subdivided for anterograde (IS: n=32; ID: n=26), reversing (IS: n=21; ID: n=7), and retrogade (IS: n=26; ID n=16) directions. Total percent time in motion for all moving mitochondria regardless of direction (excluding pauses) was calculated and subdivided for anterograde, reversing and retrograde (IS: n=98; ID: n=70) directions. The percent time mitochondria spend pausing was also calculated and plotted (IS: n=98; ID: n=70). All scatter plots display the median with the interquartile range from data pooled from two unique neuron culture preparations. An unpaired Mann-Whitney test was performed for all mitochondrial motility, density, and population parameters to compare the medians of the non-parametric datasets with alpha threshold = 0.05. Statistically significant discoveries (*) were determined after correction with FDR analysis using Benjamini-Hochberg method (q=5%).

**Table 1:** Definitions of mitochondrial motility parameters.

| Parameter | Definition | Figure |
| --- | --- | --- |
| **Mitochondrial Densities** |  |  |
| Anterograde density | # of mitochondria with only anterograde motion, with or without pausing, per $\mu\text{m}$ of axon | 3C |
| Reversal density | # of mitochondria with at least one change in motion directionality per $\mu\text{m}$ of axon | 3C |
| Retrograde density | # of mitochondria with only retrograde motion, with or without pausing, per $\mu\text{m}$ of axon | 3C |
| Moving density | Sum of # of moving mitochondria per $\mu\text{m}$ of axon for all motions (anterograde, reversal, and retrograde) | ---- |
| Stationary density | Sum of # of stationary mitochondria per $\mu\text{m}$ of axon | ---- |
| Total density | Total number of mitochondria per $\mu\text{m}$ axon (moving and stationary mitochondria) | 3C |
| **Mitochondrial Speed and Velocity** |  |  |
| Average Speed | Average speed of individual moving mitochondria including pauses and all directionalities (anterograde, reversal, and retrograde) | 3A |
| Net velocity | Average net velocity of individual moving mitochondria with net motion irrespective of directionality | 3A |
| Net anterograde velocity | Average net velocity of individual moving mitochondria with net motion in the anterograde direction | 3A |
| Net retrograde velocity | Average net velocity of individual moving mitochondria with net motion in the retrograde direction | 3A |
| Average segmental speed | Average of all segmental speeds (anterograde and retrograde) per mitochondrion regardless of direction | 3B |
| Anterograde segmental velocity | Average velocity of combined anterograde segments | 3B |
| Retrograde segmental velocity | Average velocity of combined retrograde segments | 3B |
| **Mitochondrial Run Lengths** |  |  |
| Average run length | Average length of combined segmental runs per mitochondrion regardless of direction | ---- |
| Anterograde run length | Average length of segmental run in the anterograde direction | ---- |
| Retrograde run length | Average length of segmental run in the retrograde direction | ---- |
| **Mitochondrial Motion and Pausing** |  |  |
| Percent time moving | Total percent time individual mitochondria are in motion (anterograde or retrograde), excluding pauses | 3D |
| Percent time anterograde motion | Total percent time individual moving mitochondria spend in anterograde direction | 3D |
| Percent time retrograde motion | Total percent time individual moving mitochondria spend in retrograde direction | 3D |
| Percent time pausing | Total percent time individual mitochondria spend pausing | 3D |
| Total pause frequency | Average number of pauses per individual moving mitochondria regardless of movement direction preceding or following the pause | 3D |
| Anterograde pause frequency | # of pauses per individual mitochondria between movements in the anterograde direction | 3D |
| Retrograde pause frequency | # of pauses per individual mitochondria between movements in the retrograde direction | 3D |
| Reversal pause frequency | # of pauses per individual mitochondria between reversal movements | 3D |
| Total pause duration | Average duration of all pauses for a given moving mitochondrion regardless of movement direction preceding or following the pause | 3D |
| Anterograde pause duration | Average duration of pauses for individual mitochondria with anterograde motion preceding and following the pause | 3D |
| Retrograde pause duration | Average duration of pauses for individual mitochondria with retrograde motion preceding and following the pause | 3D |
| Reversal pause duration | Average duration of pauses for individual mitochondria with a change in direction following the pause | 3D |

### Iron chelation during axonogenesis impairs axonal growth and branching

To assess if ID affects early axon growth and branching, semi-automated tracing of the entire axonal arbor from individual neurons was performed (Figure 5A). Sholl analysis of axonal tracings revealed significantly decreased (p≤0.05) number of axonal crossings between 440µm-850µm from the soma in iron-deficient compared to iron-sufficient hippocampal neurons, suggesting impaired axonal growth and/or branching (Figure 5B). To determine whether ID specifically impaired axonal outgrowth or *de novo* axon branching, the number and length of axonal processes were quantified. Consistent with the Sholl analysis, the primary (p<0.001) and total length (p=0.017) of the axonal arbor (primary segment + branches) was reduced in iron-deficient neurons (Figure 5B, C). ID did not alter the number of branches or the total branch length (Figure 6B, C); however, ID significantly reduced the average branch length (p=0.001) compared to IS neurons (Figure 6A), suggesting impaired axon linear growth but not branch formation/retraction. To further explore this, a MANOVA was performed using statistical tests by Wilk’s lambda and Pillai’s trace (Supplemental Table 1). Data are tabulated (Supplemental Table 1) ranking each modeled multivariate interaction by F-value (highest to lowest) with its corresponding raw and FDR-adjusted p-value and effect size (ŋ^2^ from Wilk’s and Pillai’s). Significant differences between IS and ID for all interactions were reported from the MANOVA except for interaction between intersection and total branch length (p = 0.051) and intersections and number of branches (p = 0.087), (Supplemental Table 1). The strongest interaction was seen between radius and primary axon length (p < 0.001). Significant group-dependent differences in covariance between number of branches and primary axon length (p = 0.008) and number of branches and average branch length (p = 0.017) was captured in the multivariate analysis.

**Figure 5:**
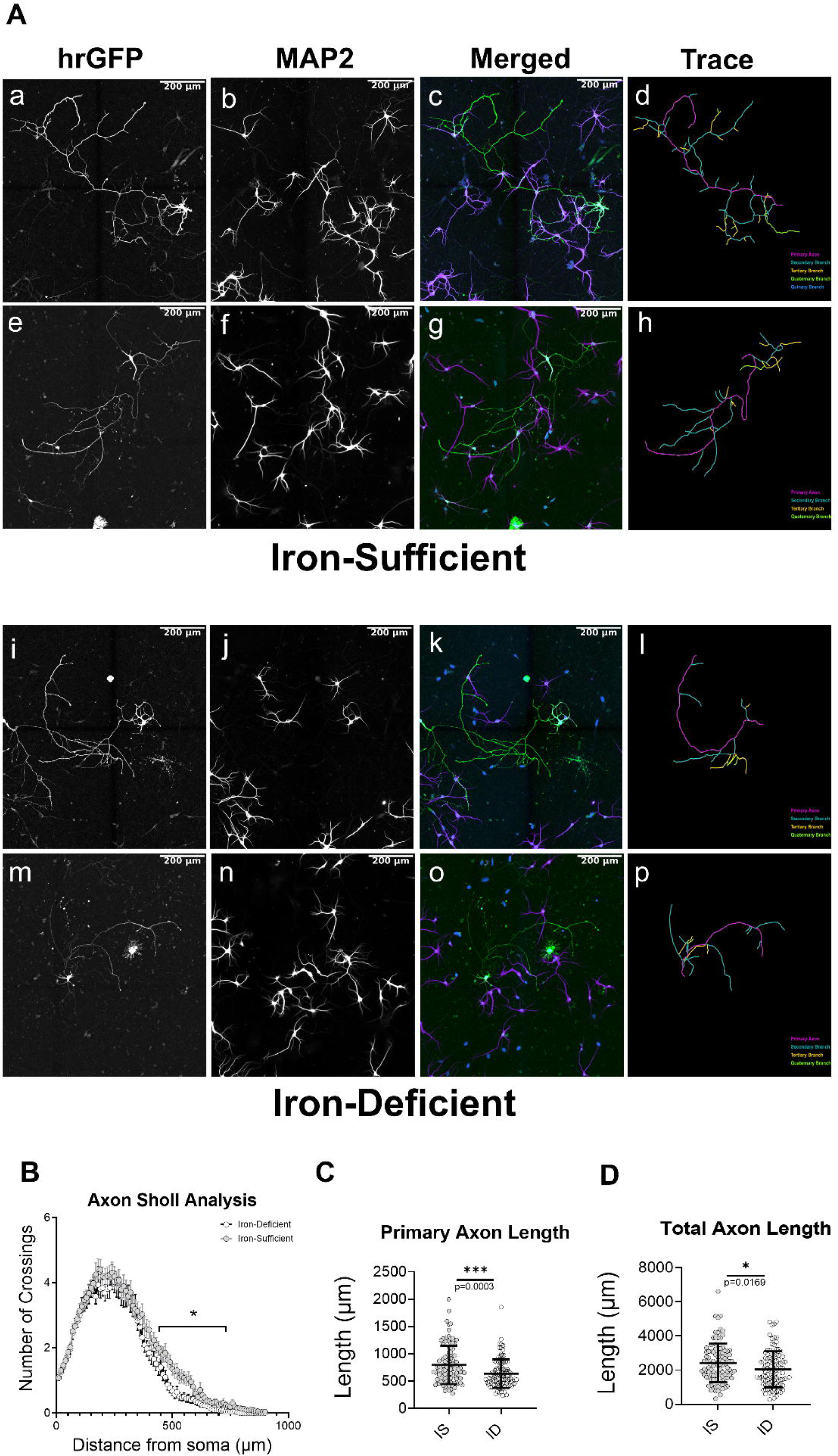
ID impairs hippocampal neuron axonal growth. Hippocampal neuronal cultures isolated from E16 mice were nucleofected with an hrGFP-expressing plasmid at time of plating. Cultures were then treated with 5-FU and with or without DFO at 3 DIV after axon specification. At 7 DIV, the neuronal cultures were fixed and processed for immunocytochemistry against MAP2 (dendrite specific marker). Individual hrGFP-positive neurons were imaged using a Zeiss Celldiscoverer 7 and Zeiss Axiocam 506 mono camera with a 20x (NA 0.7) objective and 0.5x magnification. The hrGFP-positive and MAP2-negative axon was distinguished from MAP2/hrGFP double-positive dendrites to complete axonal tracing using FIJI and Simple Neurite Tracer. **A)** Two representative grayscale images are shown of iron-sufficient and -deficient axons with corresponding hrGFP (neuron filler), MAP2 (dendrites), merged hrGFP/MAP2/DAPI channels, and skeletonized tracing. **B)** Sholl-curve (mean±SEM; n=100 neurons). An asterisk (*) indicates statistical significance at a given distance from the soma by repeated measures 2-way ANOVA and Tukey’s post-hoc test. Univariate morphological measurements were calculated from simple neurite tracer analysis of **C)** primary axon length and **D)** total axon length. Data points representing individual neurons (n=100) are displayed with Mean ± SD. An asterisk (*) indicates statistical significance via Welch’s t-test (p-values ≤ 0.05).

**Figure 6:**
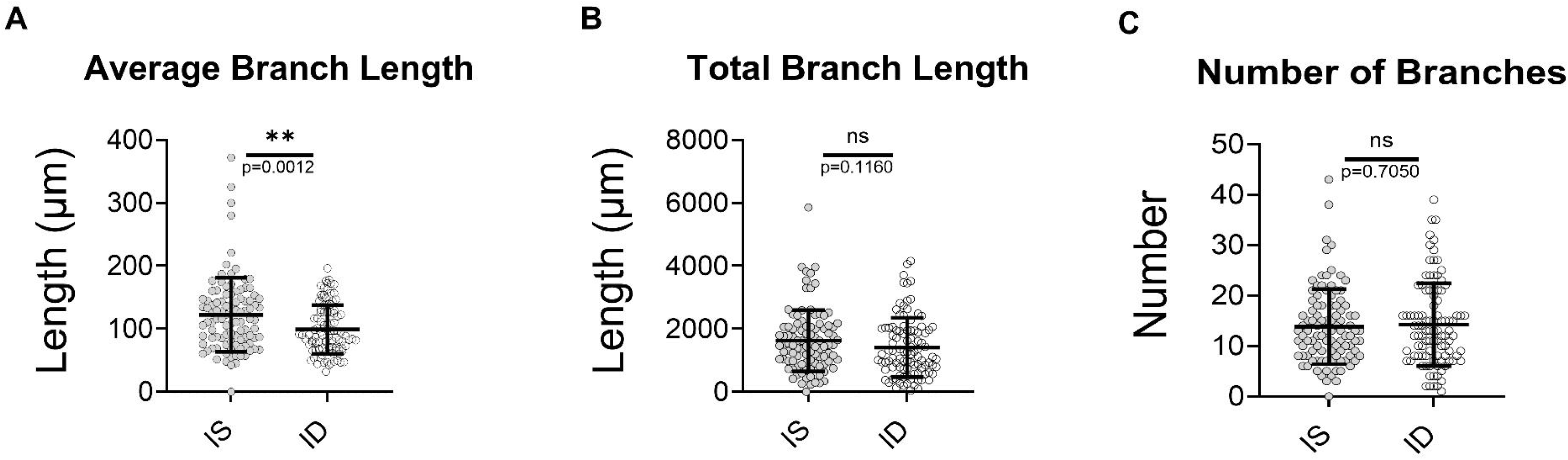
Iron deficiency impairs hippocampal neuron axonal branch growth but not formation. Axonal traces (see Fig 5) were used to measure and calculate **A)** average branch length, **B)** total branch length, and **C)** number of branches per axon. Data points representing individual neurons (n=100) are displayed with Mean ± SD. An asterisk (*) indicates statistical significance via Welch’s t-test (p-values ≤ 0.05).

Pairwise interactions are displayed as a pairplot with overlaid correlation values (R^2^) for IS and ID neurons (Supplemental Figure 1). Comparisons for regressions statistics between IS and ID neurons for each pairwise interactions was also assessed and displayed as a heatmap scaled by FDR-adjusted p-values but no differences were observed (Supplemental Figure 2). A corresponding table (Supplemental Table 2) displays the resulting linear regression statistics for each pairwise comparison (Supplemental Figure 2), listing each x-and y-variable pair, effect size, upper and lower confidence intervals, R^2^ values, raw p-values and FDR adjusted p-values comparing between IS and ID neurons (Supplemental Table 2). The resulting linear regression statistics for each pairwise comparison (Supplemental Figure 1), was also tabulated for each x-and y-variable pair, slope, intercept, R^2^ values, raw p-values and FDR adjusted p-values and listed for IS and ID neurons, independently (Supplemental Table 3). The strongest pair association with significant differences was shown between branch length and total axon length (IS: R^2^ = 0.910, p < 0.001, ID: R^2^ = 0.948, p < 0.001) and total branch length and number of branches (IS: R^2^ = 0.564, p < 0.001, ID: R^2^ = 0.665, p < 0.001) in both ID and IS neurons. Pairwise association between number of branches and primary axon length was insignificant and not strongly correlated for both IS and ID neurons (IS: R^2^ =0.625, p = 0.772, ID: R^2^ = 0.198, p = 0.441).

### Chronic ID reduces PSD95 puncta density

Neuronal ID causes deficits in both axonal (Figure 5, 6) and dendritic (Bastian et al., 2016, 2017, 2022) growth. We hypothesized that this will affect the long-term ability of iron-deficient neurons to form synapses. To test this, we electroporated E15.5 hippocampal cells with a plasmid expressing an eGFP-tagged intrabody against endogenous PSD95 (Gross et al 2013) and then allowed iron-sufficient and -deficient neurons to grow to 18DIV, a period of rapid synaptogenesis (Figure 7A). 18DIV iron-deficient neurons displayed a 30% decrease (p = 0.003) in dendritic PSD95 puncta density compared to iron-suffient neurons (Figure 7B), indicating a reduction in the potential of iron-deficient neurons to form synapses.

**Figure 7:**
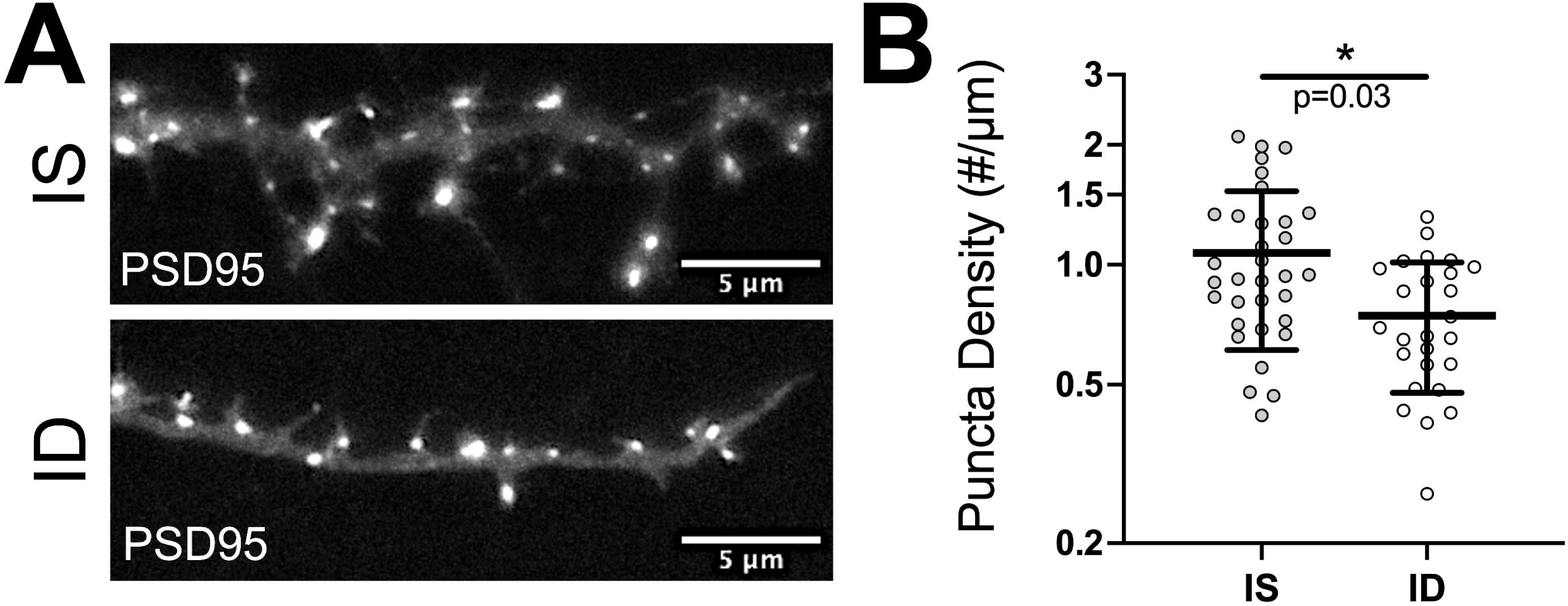
Chronic iron deficiency reduces dendritic PSD95 puncta density. **A)** Representative 18DIV images of fluorescently labeled post-synaptic puncta in dendrites from hippocampal neurons transfected prior to plating with PSD95.FingR-GFP (labels endogenous PSD95; post-synaptic marker) for iron-sufficient (top panel) and iron-deficient (bottom panel) groups. **B)** Quantification of dendritic PSD95 puncta density (iron-sufficient: n=38 neurons; iron-deficient: n=33 neurons). Data points representing individual neurons (n=100) are displayed with Mean ± SD. An asterisk (*) indicates statistical significance via unpaired t-test (p ≤ 0.05).

## Discussion

Optimal brain circuit formation depends on proper establishment of neuronal polarity, a process that is specific and unique to neurons, and includes both axonogenesis and dendritogenesis (M. Zhang et al., 2019). Early-life ID impairs neurocircuit formation and function, particularly in circuits involving hippocampal pyramidal neurons (e.g., learning and memory) (Brunette et al., 2010; Carlson et al., 2010; Fretham et al., 2011, 2012; Georgieff et al., 2018; Jorgenson et al., 2003, 2005; Radlowski & Johnson, 2013). Extensive prior work has shown hippocampal neuron dendrite growth and branching deficits during early-life ID (Bastian et al., 2016, 2022; Brunette et al., 2010; Carlson et al., 2010; Fretham et al., 2012; Jorgenson et al., 2003). A simplified dendritic arbor provides a post-synaptic explanation for the neurocircuit behavioral impairments caused by fetal-neonatal ID. However, the effects of ID on the pre-synaptic, axonal side (outside of myelination deficits (Lozoff et al., 2006; Todorich et al., 2009)) remain largely unknown. Axonogenesis is an energetically demanding process and thus energy production deficits often cause morphological aberrations in axonal development (Harris & Attwell, 2012). In the current study, neuronal ID reduced basal and ATP-linked mitochondrial respiration, ATP production rate, and neuronal ATP concentrations, suggesting a whole neuron level reduction in energy capacity of ID hippocampal neurons at 7DIV, during this period of rapid axon growth and branching. This is consistent with previous observations of impaired mitochondrial energetic capacity in the same neuron culture model, during the period of rapid dendritic growth and arborization (11-18DIV) (Bastian et al., 2016, 2019, 2022). This finding suggests that impaired iron-dependent mitochondrial function may also lead to axonal structural deficits, prior to the onset of rapid dendrite growth and branching.

In our primary embryonic mouse hippocampal neuron culture model of fetal-neonatal ID, iron chelation begins after axon initiation, which allows us to examine effects on axonal growth and branching without potential confounding effects on initial axon formation. ID caused gross axon morphological deficits with a reduction in overall axon complexity at 440µm-850µm from the soma, which was caused by a reduction in axon and branch length but not branch number. Multivariate analysis revealed group-dependent differences in covariance between primary axon length to number of branches and the average branch length to number of branches not captured in the univariate data. This would indicate that while the mean number of branches are not affected by the effects of ID at this stage of axonal development, the stochiometric relationship between axon or branch length and number of branches is altered by ID (i.e., length of axons does not correlate as closely with the number of branches). However, within group pairwise regression statistic comparisons for variable interactions between length variables (primary axon and average branch length) and number of branches revealed no significant differences. Thus, our findings indicate that iron is critical for axonal growth and elongation but not for branch number and that this is likely driven through impaired iron-dependent mitochondrial energetics during early axonogenesis (Figure 8).

**Figure 8:**
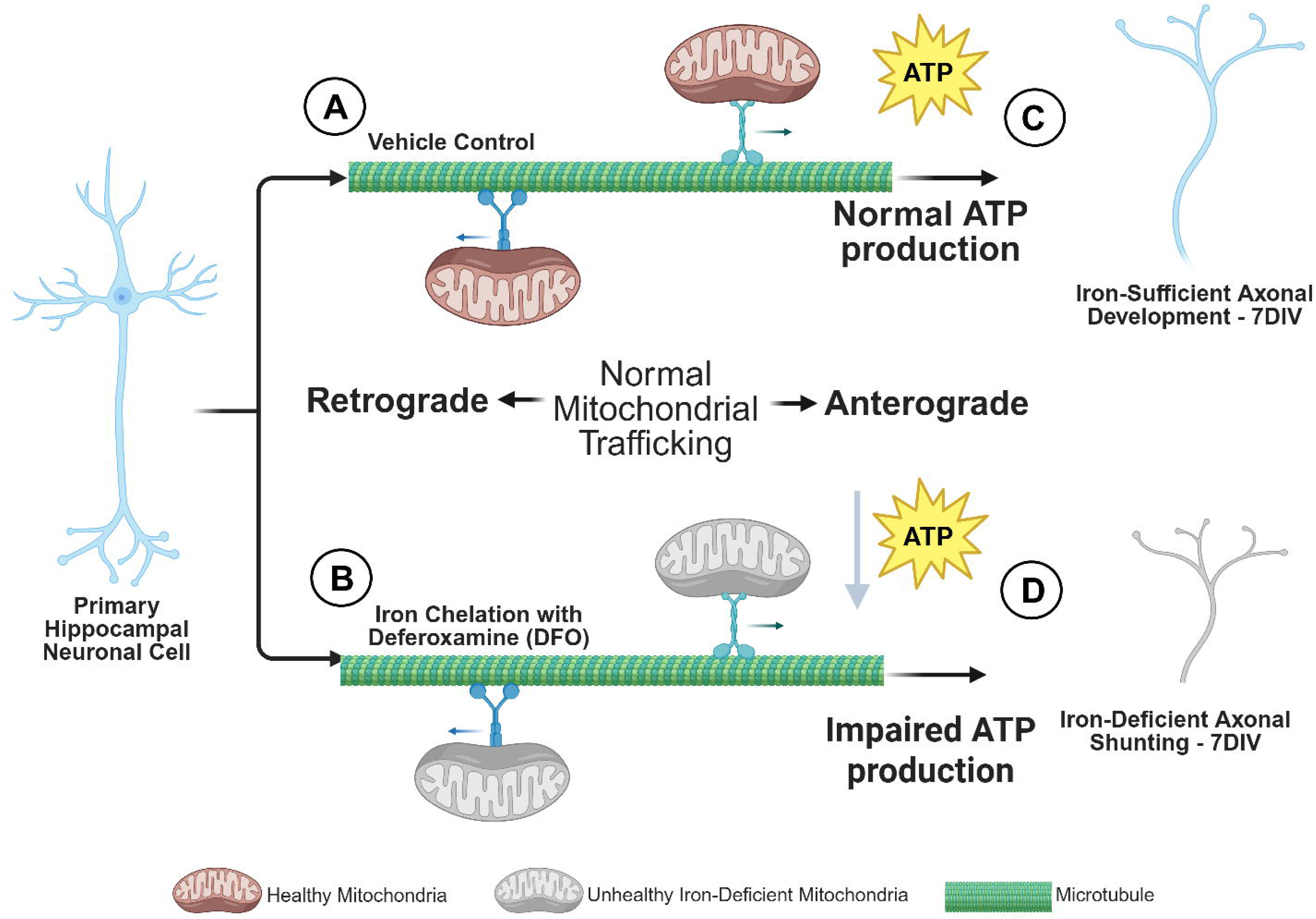
Summary of ID effects on axonal mitochondria during axonogenesis: Representative illustration of **A)** iron-sufficient (Vehicle Control) and **B)** iron-deficient (DFO chelated) hippocampal neuron at 3 DIV. **C)** Iron-sufficient hippocampal axon is modeled with healthy mitochondria (brown) exhibiting normal mitochondrial trafficking along microtubule (green) in retrograde and anterograde direction with normal ATP production by 7DIV. Axonal development outcome in primary and branching length is normal (blue). **D)** DFO treated hippocampal neurons modeled with unhealthy iron-deficient mitochondria (gray) exhibit normal trafficking along microtubule (green) in retrograde and anterograde direction but impaired ATP production by 7DIV. Resulting neuronal phenotype (gray) displays shunted primary and branching axonal length at 7DIV with no change in number of branches. Axonal mitochondrial trafficking and branch number appears to be prioritized in ID neurons over energy production and axon growth. The mitochondrial density in the axonal compartment at 7DIV was also unaltered in iron-deficient neurons.

A large body of literature has shown that fetal-neonatal ID impairs neonatal dendritic arborization (a largely postnatal process in rodents), causing long-term deficits in dendrite structure and function (Bastian et al., 2016, 2022; Brunette et al., 2010; Carlson et al., 2010; Fretham et al., 2012; Greminger & Mayer-Pröschel, 2015; Jorgenson et al., 2003; Perng et al., 2021; Figure 7). Given that axonogenesis precedes dendritogenesis by several days in rodents and weeks in humans (Semple et al., 2013), ID-induced stunting of the primary axon and its branches demonstrates that the proper structural development of neurons requires sufficient iron much earlier than previously thought. This has implications for axons facing the challenge of reaching distant targets at the proper time during embryonic/neonatal brain development. Since axonal pre-synaptic activity is important for proper neurogenesis, cell migration and dendrite outgrowth, branching, and spine/post-synaptic density formation (Andreae & Burrone, 2014; Ghosh & Shatz, 1992; O’Leary & Koester, 1993; Paus, 2023; Yamashita, Aoki, et al., 2016; Yamashita, Yamane, et al., 2016), our data suggest that the ID-induced early axonal growth deficit may be an important contributing factor to the known long-term dendrite, synaptic, and neurocircuit consequences of fetal-neonatal ID. Whether this decrease in axonal complexity in ID hippocampal neurons occurs during *in vivo* early axonogenesis is an important unanswered question that has critical implications for overall developmental circuitry deficits and long-term propagation dynamics for the mature formerly iron-deficient brain (Ofer et al., 2017).

Since anterograde trafficking of cargo is crucial to meet the demands of developing neuronal processes (Millecamps & Julien, 2013), decreased mitochondrial motility was hypothesized to contribute to the stunted axonal growth due to ID. Consistent with unchanged dendrite mitochondrial density at 11 DIV (Bastian et al., 2019), mitochondrial density was unaltered in the terminal (50-100µm from beginning of growth cone) of 7 DIV iron-deficient axons. Previous work with our hippocampal neuron culture model of ID has shown that at 11 DIV, ID reduces average mitochondrial speed due to increased pause frequency and decreases anterograde motion of moving mitochondria to energy-demanding terminal processes of developing dendrites, ultimately leading to reduced mitochondrial density in terminal dendrites at 18DIV (Bastian et al., 2019). However, during hippocampal neuron axonogenesis, significant differences in mitochondrial motility were not observed between IS and ID neurons. Given ID effects on mitochondrial energy production during axon growth and the known role of ATP in axonal mitochondrial motility (Bastian, 2019; Lewis et al., 2013; Sheng, 2014, 2017), this finding was unexpected. These data suggest that axonal mitochondria trafficking is prioritized over other ATP-dependent processes and indicates that there are potential compensatory mechanisms to maintain trafficking during neuronal ID (Cheng et al., 2022; Sheng, 2014; Wang et al., 2021). Overall, these findings imply that the developmental impact of ID on axon growth is due to impaired mitochondrial bioenergetics and ATP production but not reduced mitochondrial trafficking to the axonal shaft and growth cone (Figure 7).

Although iron’s role in mitochondrial energetics is critical to cellular health and development, iron is necessary for many additional important cellular processes that could contribute to impaired axon development during ID such as regulation of gene transcription, neurotransmitter synthesis, and myelination (Hare et al., 2013). The reduced non-mitochondrial respiration observed in iron-deficient neurons likely indicates reduced activity of other iron-dependent oxygen consuming enzymes (e.g., Tet dioxygenases, lysine histone demethylases (Kdms), prolyl hydroxylase, desaturases, etc.). Tet and Kdm enzymes catalyze the demethylation of DNA and histones, respectively, and are responsible for iron-dependent epigenetic regulation of transcription for key neurodevelopmental genes (e.g., Bdnf) (Barks et al., 2018; Tran et al., 2015). In addition, the iron-dependent enzyme, prolyl hydroxylase, is necessary for the regulation of hypoxia inducible factor 1 α (HIF1α), a regulator of many metabolic and neurodevelopmental genes (Hare et al., 2013; Xu et al., 2017; D. L. Zhang et al., 2014). Iron-containing desaturase enzymes are responsible for the production of mono- and poly-unsaturated fatty acids, key metabolic substrates and components of neuronal membranes (Bazinet & Layé, 2014). Future work is needed to explore the prioritization of iron to these enzymes during neuronal ID and their relative contribution to iron-dependent neuron structural development.

An important unanswered question is why ID-induced mitochondrial energy deficits impair axonal growth but not branch number (Figure 6). Tao *et al* showed mitochondrial ATP dependent branch formation in immature hippocampal neurons with stationary mitochondria present at branch points (Tao et al., 2014). In early sensory axons, impairment of neural growth factor (NGF) mediated mitochondrial respiration impaired actin but not microtubule polymerization indicating NGF mediated axonal branching requires local ATP regulation of actin dynamics (Smith & Gallo, 2018; Spillane et al., 2013). Since we examined only whole cell mitochondrial respiration and ATP production, it is unclear whether the local ATP concentration near axon branch points was maintained or not. Since it is known that stationary mitochondria along neuronal processes are important for branch formation, it is possible that *de novo* axonal branching does not necessarily depend on mitochondrial ATP production but rather other mitochondrial activities (e.g. calcium buffering). Since we did not observe a change in axonal mitochondrial motility or density, one explanation is that branch number is maintained because ID does not alter mitochondrial localization near branch points. Future studies with temporal and spatial resolution of ATP dynamics targeted to subcellular regions of interest (e.g., branch points and growth cone) are necessary to better understand the relationship between mitochondrial energetics/trafficking and axonal growth/branching during ID (Marvin et al., 2024). In addition, many other mechanisms (e.g., mRNA trafficking, actin, and microtubule polymerization, and recruitment of other organelles) are also involved in determining branch formation (Lewis et al., 2013; Smith & Gallo, 2018) and will need to be explored during neuronal ID.

Most pre-clinical studies on the effects of fetal-neonatal ID on brain development have focused on postnatal neurodevelopmental processes (e.g., dendritic arborization, myelination, and synapse formation). ID-induced axon structural deficits have implications for the proper neurocircuit sculpting of the brain starting during embryonic development. Translationally, these findings provide biological plausibility for shifting the focus from treating iron-deficient children to monitoring and management of maternal iron status throughout pregnancy (Georgieff et al., 2018). Indeed, several clinical studies demonstrate that maternal iron status during pregnancy predicts the risk of offspring developing autism spectrum, attention deficit hyperactive, and schizophrenia disorders (reviewed in (Georgieff et al., 2018; Maxwell & Rao, 2022); all thought to be disorders of circuit construction. This is further supported by studies demonstrating that gestational ID alters the balance of excitatory to inhibitory neurons in the embryonic brain (Rudy et al., 2023). The long-term implications of impaired iron-dependent neurodevelopment are further underscored by studies showing that early-life ID causes Parkinsonian-like neurodegeneration (Matak et al., 2016), highlighting the need for early interventions. More efforts have been focusing on using and developing mitochondrial targeted therapies to treat neurodegenerative and neuropsychiatric disorders, thus showing a converging interest at optimizing mitochondrial health (Andreyev et al., 2024; Callegari et al., 2025; Włodarczyk et al., 2018). Improved understanding of the interface between ID and mitochondrial function in early-life neuronal development could bring about preventative solutions and interventions to optimize brain health for populations at risk of ID as well as those suffering from seemingly unrelated brain-health disorders with potential origins in iron or mitochondrial dysregulation. Altogether, our observations underscore the need to devote future efforts to understanding the effects of ID (and other common nutrient disruptions) on embryonic brain development processes to understand the stage of development that interventions are required to prevent the long-lasting neurobehavioral deficits caused by fetal-neonatal ID.

---

**Movie 1: Representative movies of IS and ID mitochondrial motility in terminal axon processes.** Movies show time-lapse imaging of mCherry-Mito-7 positive mitochondria in 7 DIV **A)** IS and **B)** ID axonal terminal processes from E16 hippocampal neuronal cultures treated with 5-FU and DFO at 3 DIV. The mitochondria for this representative movie were imaged every 1s for 1.7 min and movies are set to 30 frames per second. Images were straightened using the Fiji “Straightened” function to better visualize the region of interest in the axon terminal. Scale bars represent 5µm.

## Author Contributions

D.C.M., K.D., and T.R.M. and T.W.B wrote the first draft of the paper; D.C.M., T.W.B, M.K.G., and L.M.L. designed the experiments and edited the paper; D.C.M., K.D., L.H.C., D.J.M. and T.W.B. performed the experiments; D.C.M., K.D., T.W.B. and T.R.M. analyzed the data; The paper was read and approved by all authors before submission.

## Funding Sources and Disclosures

The work from these studies was supported by the National Institute of Health (NIH) Grant R01HD094809 to M.K.G., NIH R21HD106043 to T.W.B. and the Howard Hughes Medical Institute (HHMI) Gilliam Fellows program award GT15879 to T.W.B. and D.C.M. T.R.M. was supported by an NIH T32 training grant (T32HL007062).

## Conflict of Interest

The authors declare no conflict of interest to report.

## Code and Data Availability

Code and data are publicly available through the Data Repository for the University of Minnesota (DRUM) at https://hdl.handle.net/11299/166578

## Acknowledgements

With utmost gratefulness, we acknowledge the support of other lab members of the Bastian, Georgieff and Lanier laboratories for their assistance with preparation of cell cultures, troubleshooting and valuable feedback.

## Notes

### Competing Interest Statement

The authors have declared no competing interest.

### Summary of Updates

New data were added showing chronic iron deficiency effects on dendritic PSD95 puncta density. Additional writing and references in the Introduction and Discussion was added.

## References

Andreae L.C., Burrone J. (2014) The role of neuronal activity and transmitter release on synapse formation. Current Opinion in Neurobiology 27, 47–52. 10.1016/j.conb.2014.02.008

Andreyev, A. Y., Yang, H., Doulias, P. T., Dolatabadi, N., Zhang, X., Luevanos, M., Blanco, M., Baal, C., Putra, I., Nakamura, T., Ischiropoulos, H., Tannenbaum, S. R., & Lipton, S. A. (2024). Metabolic Bypass Rescues Aberrant S-nitrosylation-Induced TCA Cycle Inhibition and Synapse Loss in Alzheimer’s Disease Human Neurons. Advanced Science, 11(12). 10.1002/advs.202306469

Appolus EE, & Okoli CN. (2022). A Robust Comparison Powers of Four Multivariate Analysis of Variance Tests. European Journal of Statistics and Probability, 10(1), 11–20. 10.37745/ejsp.2013

Barks, A., Fretham, S. J. B., Georgieff, M. K., & Tran, P. V. (2018). Early-Life Neuronal-Specific Iron Deficiency Alters the Adult Mouse Hippocampal Transcriptome. Journal of Nutrition, 148(10), 1521–1528. 10.1093/jn/nxy125

Bastian, T. W. (2019). Potential mechanisms driving mitochondrial motility impairments in developing iron-deficient neurons. Journal of Experimental Neuroscience, 13. 10.1177/1179069519858351

Bastian, T. W., Rao, R., Tran, P. V., & Georgieff, M. K. (2020). The Effects of Early-Life Iron Deficiency on Brain Energy Metabolism. In Neuroscience Insights (Vol. 15). SAGE Publications Ltd. 10.1177/2633105520935104

Bastian, T. W., von Hohenberg, W. C., Georgieff, M. K., & Lanier, L. M. (2019). Chronic energy depletion due to iron deficiency impairs dendritic mitochondrial motility during hippocampal neuron development. Journal of Neuroscience, 39(5), 802–813. 10.1523/JNEUROSCI.1504-18.2018

Bastian, T. W., Von Hohenberg, W. C., Kaus, O. R., Lanier, L. M., & Georgieff, M. K. (2022). Choline Supplementation Partially Restores Dendrite Structural Complexity in Developing Iron-Deficient Mouse Hippocampal Neurons. Journal of Nutrition, 152(3), 747–757. 10.1093/jn/nxab429

Bastian, T. W., Von Hohenberg, W. C., Mickelson, D. J., Lanier, L. M., & Georgieff, M. K. (2016). Iron Deficiency Impairs Developing Hippocampal Neuron Gene Expression, Energy Metabolism, and Dendrite Complexity. Developmental Neuroscience, 38(4), 264–276. 10.1159/000448514

Bazinet, R. P., & Layé, S. (2014). Polyunsaturated fatty acids and their metabolites in brain function and disease. In Nature Reviews Neuroscience (Vol. 15, Issue 12, pp. 771–785). Nature Publishing Group. 10.1038/nrn3820

Brunette, K. E., Tran, P. V., Wobken, J. D., Carlson, E. S., & Georgieff, M. K. (2010). Gestational and neonatal iron deficiency alters apical dendrite structure of CA1 pyramidal neurons in adult rat hippocampus. Developmental Neuroscience, 32(3), 238–248. 10.1159/000314341

Callegari, S., Kirk, N. S., Gan, Z. Y., Dite, T., Cobbold, S. A., Leis, A., Dagley, L. F., Glukhova, A., & Komander, D. (2025). Structure of human PINK1 at a mitochondrial TOM-VDAC array. Science. 10.1126/science.adu6445

Carlson, E. S., Fretham, S. J. B., Unger, E., O’Connor, M., Petryk, A., Schallert, T., Rao, R., Tkac, I., & Georgieff, M. K. (2010). Hippocampus specific iron deficiency alters competition and cooperation between developing memory systems. Journal of Neurodevelopmental Disorders, 2(3), 133–143. 10.1007/s11689-010-9049-0

Carlson, E. S., Stead, J. D. H., Neal, C. R., Petryk, A., & Georgieff, M. K. (2007). Perinatal iron deficiency results in altered developmental expression of genes mediating energy metabolism and neuronal morphogenesis in hippocampus. Hippocampus, 17(8), 679–691. 10.1002/hipo.20307

Carlson, E. S., Tkac, I., Magid, R., O’Connor, M. B., Andrews, N. C., Schallert, T., Gunshin, H., Georgieff, M., & Petryk, A. (2009). Iron is essential for neuron development and memory function in mouse hippocampus 1-3. Journal of Nutrition, 139(4), 672–679. 10.3945/jn.108.096354

Cheng, X. T., Huang, N., & Sheng, Z. H. (2022). Programming axonal mitochondrial maintenance and bioenergetics in neurodegeneration and regeneration. In Neuron (Vol. 110, Issue 12, pp. 1899–1923). Cell Press. 10.1016/j.neuron.2022.03.015

Cheng, X. T., & Sheng, Z. H. (2021). Developmental regulation of microtubule-based trafficking and anchoring of axonal mitochondria in health and diseases. In Developmental Neurobiology (Vol. 81, Issue 3, pp. 284–299). John Wiley and Sons Inc. 10.1002/dneu.22748

Deungria, M., Rao, R., Wobken, J. D., Luciana, M., Nelson, C. A., & Georgieff, M. K. (2000). Perinatal Iron Deficiency Decreases Cytochrome c Oxidase (CytOx) Activity in Selected Regions of Neonatal Rat Brain. In Pediatric Research. 10.1203/00006450-200008000-00009

Dotti, C. G., Sullivan, C. A., & Banker, G. A. (1988). The Establishment of Polarity by Hippocampal Neurons in Culture. In The Journal of Neuroscience: Vol. I (Issue 4).

Fretham, S. J. B., Carlson, E. S., & Georgieff, M. K. (2011). The role of iron in learning and memory. Advances in Nutrition, 2(2), 112–121. 10.3945/an.110.000190

Fretham, S. J. B., Carlson, E. S., Wobken, J., Tran, P. V., Petryk, A., & Georgieff, M. K. (2012). Temporal manipulation of transferrin-receptor-1-dependent iron uptake identifies a sensitive period in mouse hippocampal neurodevelopment. Hippocampus, 22(8), 1691–1702. 10.1002/hipo.22004

Gao, Q., Zhou, Y., Chen, Y., Hu, W., Jin, W., Zhou, C., Yuan, H., Li, J., Lin, Z., & Lin, W. (2025). Role of iron in brain development, aging, and neurodegenerative diseases. Annals of Medicine, 57(1), 2472871. 10.1080/07853890.2025.2472871

Georgieff, M. K., Ramel, S. E., & Cusick, S. E. (2018). Nutritional influences on brain development. In Acta Paediatrica, International Journal of Paediatrics (Vol. 107, Issue 8, pp. 1310–1321). Blackwell Publishing Ltd. 10.1111/apa.14287

Grabrucker, A., Vaida, B., Bockmann, J., & Boeckers, T. M. (2009). Synaptogenesis of hippocampal neurons in primary cell culture. Cell and Tissue Research, 338(3), 333–341. 10.1007/s00441-009-0881-z

Greminger, A. R., & Mayer-Pröschel, M. (2015). Identifying the threshold of iron deficiency in the central nervous system of the rat by the auditory brainstem response. ASN Neuro, 7(1), 1– 10. 10.1177/1759091415569911

Gross G.G., Junge J.A., Mora R.J., Kwon H-B., Olson C.A., Takahashi T.T., Liman E.R., Ellis-Davies G.C.R., McGee A.W., Sabatini B.L., Roberts R.W., Arnold D.B. (2013) Recombinant Probes for Visualizing Endogenous Synaptic Proteins in Living Neurons. Neuron 78(6), 971–985. 10.1016/j.neuron.2013.04.017

Hare, D., Ayton, S., Bush, A., & Lei, P. (2013). A delicate balance: Iron metabolism and diseases of the brain. In Frontiers in Aging Neuroscience (Vol. 5, Issue JUL). 10.3389/fnagi.2013.00034

Harris, J. J., & Attwell, D. (2012). The energetics of CNS white matter. Journal of Neuroscience, 32(1), 356–371. 10.1523/JNEUROSCI.3430-11.2012

Insel, B. J., Schaefer, C. A., Mckeague, I. W., Susser, E. S., & Brown, A. S. (2008). Maternal Iron Deficiency and the Risk of Schizophrenia in Offspring. In Arch Gen Psychiatry (Vol. 65, Issue 10).

Jorgenson, L. A., Wobken, J. D., & Georgieff, M. K. (2003). Perinatal Iron Deficiency Alters Apical Dendritic Growth in Hippocampal CA1 Pyramidal Neurons. Developmental Neuroscience, 25(6), 412–420. 10.1159/000075667

Kim, J. W., Byun, M. S., Yi, D., Lee, J. H., Jeon, S. Y., Ko, K., Joung, H., Jung, G., Lee, J. Y., Sohn, C. H., Lee, Y. S., Kim, Y. K., & Lee, D. Y. (2021). Blood Hemoglobin, in-vivo Alzheimer Pathologies, and Cognitive Impairment: A Cross-Sectional Study. Frontiers in Aging Neuroscience, 13. 10.3389/fnagi.2021.625511

Konofal, E., Lecendreux, M., Arnulf, I., & Mouren, M.-C. (2004). Iron Deficiency in Children With Attention-Deficit/Hyperactivity Disorder.

Lewis, T. L., Courchet, J., & Polleux, F. (2013). Cellular and molecular mechanisms underlying axon formation, growth, and branching. In Journal of Cell Biology (Vol. 202, Issue 6, pp. 837–848). 10.1083/jcb.201305098

Lozoff, B., Beard, J., Connor, J., Felt, B., Georgieff, M., & Schallert, T. (2006). Long-Lasting Neural and Behavioral Effects of Iron Deficiency in Infancy.

Lozoff, B., & Georgieff, M. K. (2006). Iron Deficiency and Brain Development. Seminars in Pediatric Neurology, 13(3), 158–165.

Lozoff, B., Smith, J. B., Kaciroti, N., Clark, K. M., Guevara, S., & Jimenez, E. (2013). Functional significance of early-life iron deficiency: Outcomes at 25 years. Journal of Pediatrics, 163(5), 1260–1266. 10.1016/j.jpeds.2013.05.015

Lukowski, A. F., Koss, M., Burden, M. J., Jonides, J., Nelson, C. A., Kaciroti, N., Jimenez, E., & Lozoff, B. (2010). Iron deficiency in infancy and neurocognitive functioning at 19 years: Evidence of long-term deficits in executive function and recognition memory. Nutritional Neuroscience, 13(2), 54–70. 10.1179/147683010X12611460763689

Marvin, J. S., Kokotos, A. C., Kumar, M., Pulido, C., Tkachuk, A. N., Yao, J. S., Brown, T. A., & Ryan, T. A. (2024). iATPSnFR2: A high-dynamic-range fluorescent sensor for monitoring intracellular ATP. Proceedings of the National Academy of Sciences of the United States of America, 121(21). 10.1073/pnas.2314604121

Matak, P., Matak, A., Moustafa, S., Aryal, D. K., Benner, E. J., Wetsel, W., & Andrews, N. C. (2016). Disrupted iron homeostasis causes dopaminergic neurodegeneration in mice. Proceedings of the National Academy of Sciences of the United States of America, 113(13), 3428–3435. 10.1073/pnas.1519473113

Maxwell, A. M., & Rao, R. B. (2022). Perinatal iron deficiency as an early risk factor for schizophrenia. In Nutritional Neuroscience (Vol. 25, Issue 10, pp. 2218–2227). Taylor and Francis Ltd. 10.1080/1028415X.2021.1943996

Millecamps, S., & Julien, J. P. (2013). Axonal transport deficits and neurodegenerative diseases. In Nature Reviews Neuroscience (Vol. 14, Issue 3, pp. 161–176). 10.1038/nrn3380

Monko, T. R., Tripp, E. H., Burr, S. E., Gunderson, K. N., Lanier, L. M., Georgieff, M. K., & Bastian, T. W. (2024). Cellular Iron Deficiency Disrupts Thyroid Hormone Regulated Gene Expression in Developing Hippocampal Neurons. Journal of Nutrition, 154(1), 49–59. 10.1016/j.tjnut.2023.11.007

Monko T, Bastian T. napari-ndev: a generalizable start-to-finish napari plugin for batch bioimage processing and analysis without code [Internet]. Zenodo; 2025 [cited 2025 Feb 19]. Available from: https://zenodo.org/records/14787853

Neumann, S., Chassefeyre, R., Campbell, G. E., & Encalada, S. E. (2017). KymoAnalyzer: a software tool for the quantitative analysis of intracellular transport in neurons. Traffic, 18(1), 71–88. 10.1111/tra.12456

Ofer, N., Shefi, O., & Yaari, G. (2017). Branching morphology determines signal propagation dynamics in neurons. Scientific Reports, 7(1). 10.1038/s41598-017-09184-3

Paus, T. (2023). Tracking Development of Connectivity in the Human Brain: Axons and Dendrites. In Biological Psychiatry (Vol. 93, Issue 5, pp. 455–463). Elsevier Inc. 10.1016/j.biopsych.2022.08.019

Penrod, R. D., Campagna, J., Panneck, T., Preese, L., & Lanier, L. M. (2015). The presence of cortical neurons in striatal-cortical co-cultures alters the effects of dopamine and BDNF on medium spiny neuron dendritic development. Frontiers in Cellular Neuroscience, 9(July), 1–14. 10.3389/fncel.2015.00269

Perng, V., Li, C., Klocke, C. R., Navazesh, S. E., Pinneles, D. K., Lein, P. J., & Ji, P. (2021). Iron Deficiency and Iron Excess Differently Affect Dendritic Architecture of Pyramidal Neurons in the Hippocampus of Piglets. Journal of Nutrition, 151(1), 235–244. 10.1093/jn/nxaa326

Petry, C. D., Eaton, M. A., Wobken, J. D., Mills, M. M., Johnson, D. E., & Georgieff, M. K. (1992). Iron deficiency of liver, heart, and brain in newborn infants of diabetic mothers. 10.1016/s0022-3476(05)82554-5

Radlowski, E. C., & Johnson, R. W. (2013a). Perinatal iron deficiency and neurocognitive development. In Frontiers in Human Neuroscience (Issue SEP). Frontiers Media S. A. 10.3389/fnhum.2013.00585

Radlowski, E. C., & Johnson, R. W. (2013b). Perinatal iron deficiency and neurocognitive development. In Frontiers in Human Neuroscience (Issue SEP). Frontiers Media S. A. 10.3389/fnhum.2013.00585

Rao, R., Tkac, I., Townsend, E. L., Gruetter, R., & Georgieff, M. K. (2003). Nutritional Neurosciences Perinatal Iron Deficiency Alters the Neurochemical Profile of the Developing. In J. Nutr (Vol. 133, pp. 3215–3221). 10.1093/jn/133.10.3215

Rudy M.J., Salois G., Cubello J., Newell R., Mayer-Proschel M. (2023) Gestational iron deficiency affects the ratio between interneuron subtypes in the postnatal cerebral cortex in mice. Development 150(20), dev201068. 10.1242/dev.201068

Sahu, M. P., Nikkilä, O., Lagas, S., Kolehmainen, S., & Castrén, E. (2019). Culturing primary neurons from rat hippocampus and cortex. Neuronal Signaling, 3(2). 10.1042/NS20180207

Schindelin, J., Arganda-Carreras, I., Frise, E., Kaynig, V., Longair, M., Pietzsch, T., Preibisch, S., Rueden, C., Saalfeld, S., Schmid, B., Tinevez, J. Y., White, D. J., Hartenstein, V., Eliceiri, K., Tomancak, P., & Cardona, A. (2012). Fiji: An open-source platform for biological-image analysis. In Nature Methods (Vol. 9, Issue 7, pp. 676–682). 10.1038/nmeth.2019

Schmidt, R. J., Tancredi, D. J., Krakowiak, P., Hansen, R. L., & Ozonoff, S. (2014). Maternal intake of supplemental iron and risk of autism spectrum disorder. American Journal of Epidemiology, 180(9), 890–900. 10.1093/aje/kwu208

Semple, B. D., Blomgren, K., Gimlin, K., Ferriero, D. M., & Noble-Haeusslein, L. J. (2013). Brain development in rodents and humans: Identifying benchmarks of maturation and vulnerability to injury across species. In Progress in Neurobiology (Vols. 106–107, pp. 1–16). 10.1016/j.pneurobio.2013.04.001

Sheng, Z. H. (2014). Mitochondrial trafficking and anchoring in neurons: New insight and implications. In Journal of Cell Biology (Vol. 204, Issue 7, pp. 1087–1098). Rockefeller University Press. 10.1083/jcb.201312123

Sheng, Z. H. (2017). The Interplay of Axonal Energy Homeostasis and Mitochondrial Trafficking and Anchoring. In Trends in Cell Biology (Vol. 27, Issue 6, pp. 403–416). Elsevier Ltd. 10.1016/j.tcb.2017.01.005

Shepherd, G. M. G., & Harris, K. M. (1998). Three-Dimensional Structure and Composition of CA3 --> CA1 Axons in Rat Hippocampal Slices: Implications for Presynaptic Connectivity and Compartmentalization. Journal of Neuroscience.

Smith, G. M., & Gallo, G. (2018). The role of mitochondria in axon development and regeneration. In Developmental Neurobiology (Vol. 78, Issue 3, pp. 221–237). John Wiley and Sons Inc. 10.1002/dneu.22546

Spillane, M., Ketschek, A., Merianda, T. T., Twiss, J. L., & Gallo, G. (2013). Mitochondria Coordinate Sites of Axon Branching through Localized Intra-axonal Protein Synthesis. Cell Reports, 5(6), 1564–1575. 10.1016/j.celrep.2013.11.022

Tao, K., Matsuki, N., & Koyama, R. (2014). AMP-activated protein kinase mediates activity-dependent axon branching by recruiting mitochondria to axon. Developmental Neurobiology, 74(6), 557–573. 10.1002/dneu.22149

Todorich, B., Pasquini, J. M., Garcia, C. I., Paez, P. M., & Connor, J. R. (2009). Oligodendrocytes and myelination: The role of iron. In GLIA (Vol. 57, Issue 5, pp. 467–478). 10.1002/glia.20784

Tran, P. V, Kennedy, B. C., Lien, Y.-C., Simmons, R. A., & Georgieff, M. K. (2015). Fetal iron deficiency induces chromatin remodeling at the Bdnf locus in adult rat hippocampus. 10.1152/ajpregu.00429.2014.-Fetal

Wang, B., Huang, M., Shang, D., Yan, X., Zhao, B., & Zhang, X. (2021). Mitochondrial Behavior in Axon Degeneration and Regeneration. In Frontiers in Aging Neuroscience (Vol. 13). Frontiers Media S.A. 10.3389/fnagi.2021.650038

Włodarczyk, A., Wiglusz, M. S., & Cubała, W. J. (2018). Ketogenic diet for schizophrenia: Nutritional approach to antipsychotic treatment. Medical Hypotheses, 118, 74–77. 10.1016/j.mehy.2018.06.022

Xu, M.-M., Wang, J., & Jun-Xia, X. (2017). Regulation of iron metabolism by hypoxia-inducible factors 1 Hypoxia-inducible factors (HIFs). Acta Physiologica Sinica, 69(5), 598–610. 10.13294/j.aps.2017.0054

Yamashita, N., Aoki, R., Chen, S., Jitsuki-Takahashi, A., Ohura, S., Kamiya, H., & Goshima, Y. (2016). Voltage-gated calcium and sodium channels mediate Sema3A retrograde signaling that regulates dendritic development. Brain Research, 1631, 127–136. 10.1016/j.brainres.2015.11.034

Yamashita, N., Yamane, M., Suto, F., & Goshima, Y. (2016). TrkA mediates retrograde semaphorin 3A signaling through plexin A4 to regulate dendritic branching. Journal of Cell Science, 129(9), 1802–1814. 10.1242/jcs.184580

Zhang, D. L., Ghosh, M. C., & Rouault, T. A. (2014). The physiological functions of iron regulatory proteins in iron homeostasis - an update. In Frontiers in Pharmacology: Vol. 5 JUN. Frontiers Research Foundation. 10.3389/fphar.2014.00124

Zhang, M., Ergin, V., Lin, L., Stork, C., Chen, L., & Zheng, S. (2019). Axonogenesis Is Coordinated by Neuron-Specific Alternative Splicing Programming and Splicing Regulator PTBP2. Neuron, 101(4), 690–706.e10. 10.1016/j.neuron.2019.01.022

Zolotukhin, S., Potter, M., Hauswirth, W. W., Guy, J., & Muzyczka, N. (1996). A “Humanized” Green Fluorescent Protein cDNA Adapted for High-Level Expression in Mammalian Cells. In JOURNAL OF VIROLOGY (Vol. 70, Issue 7). https://journals.asm.org/journal/jvi

