## Supplemental for "Iron Deficiency Impairs Mitochondrial Energetics and Early Axonal Growth and Branching in Developing Hippocampal Neurons"

**Supplemental Methods:**

PSD95 Puncta Counting ImageJ Macro Code:

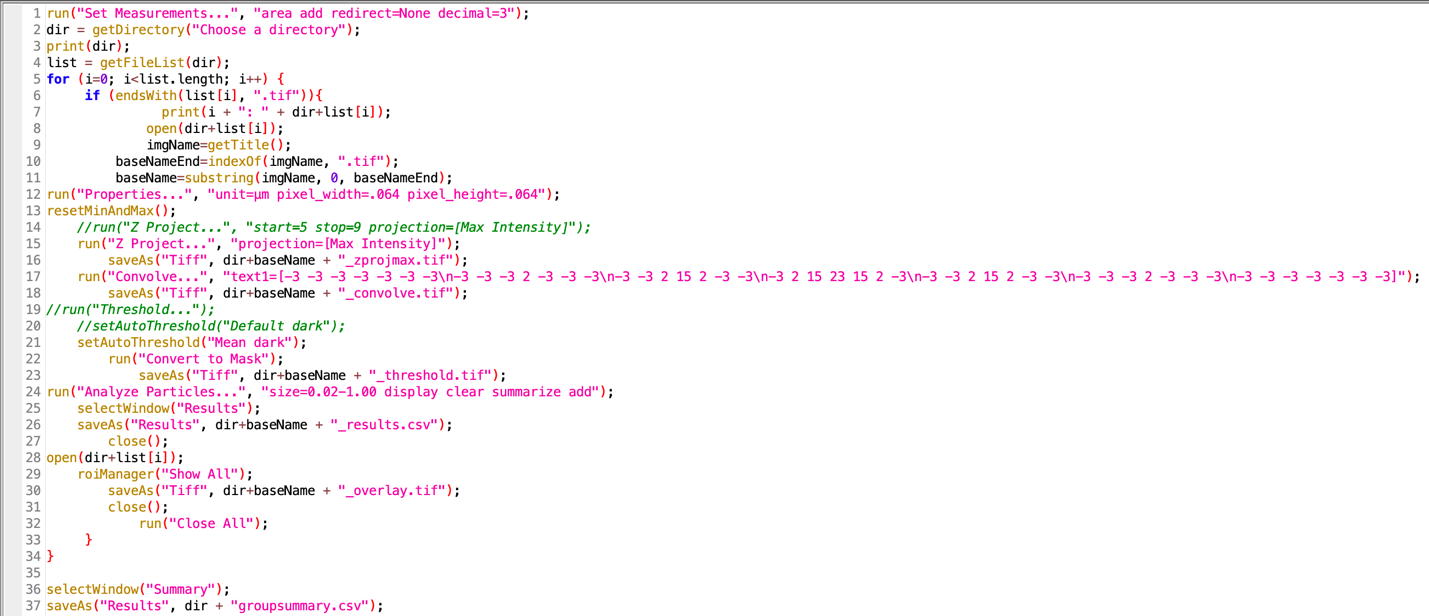

**Supplemental Data:**

**Supplemental Table 1: Multivariate analysis of axonal complexity from Sholl data.** Univariate data from Figure 4 was concatenated and used to perform multivariate analysis (n=100 neurons/group). A MANOVA was performed and the output summary table shows results from Wilk’s lambda, Pilliai’s trace, F-value (ranked highest to lowest), resulting raw and FDR adjusted p-values, and effect size (ŋ^2^) for each model interaction. Statistical significance was considered for any FDR adjusted p-value ≤ 0.05

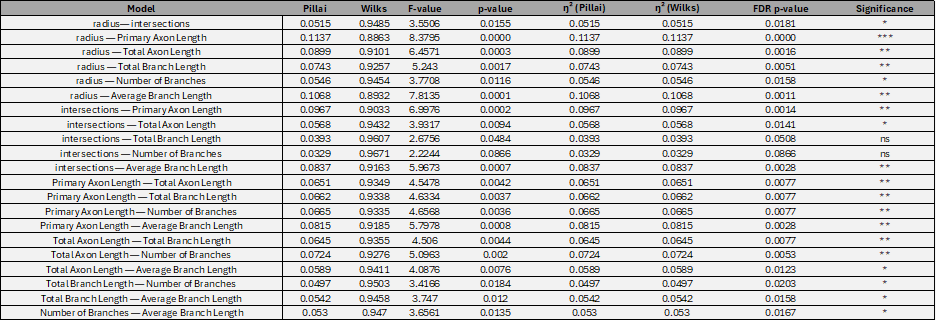

**Supplemental Figure 1: Multivariate pairplot of MANOVA Sholl data.** Multivariate scatter plots display pairwise relationships for the following variables: radius from soma (Radius), the total branch length, primary axon length, total axon length, number of branches, average branch length, and total intersections on both the y- and x-axis (n=100 neurons/group). Linear regressions for for IS (dark gray) and ID (light gray) groups are plotted with corresponding R^2^ values in each subplot to show association strength between variable pairs.

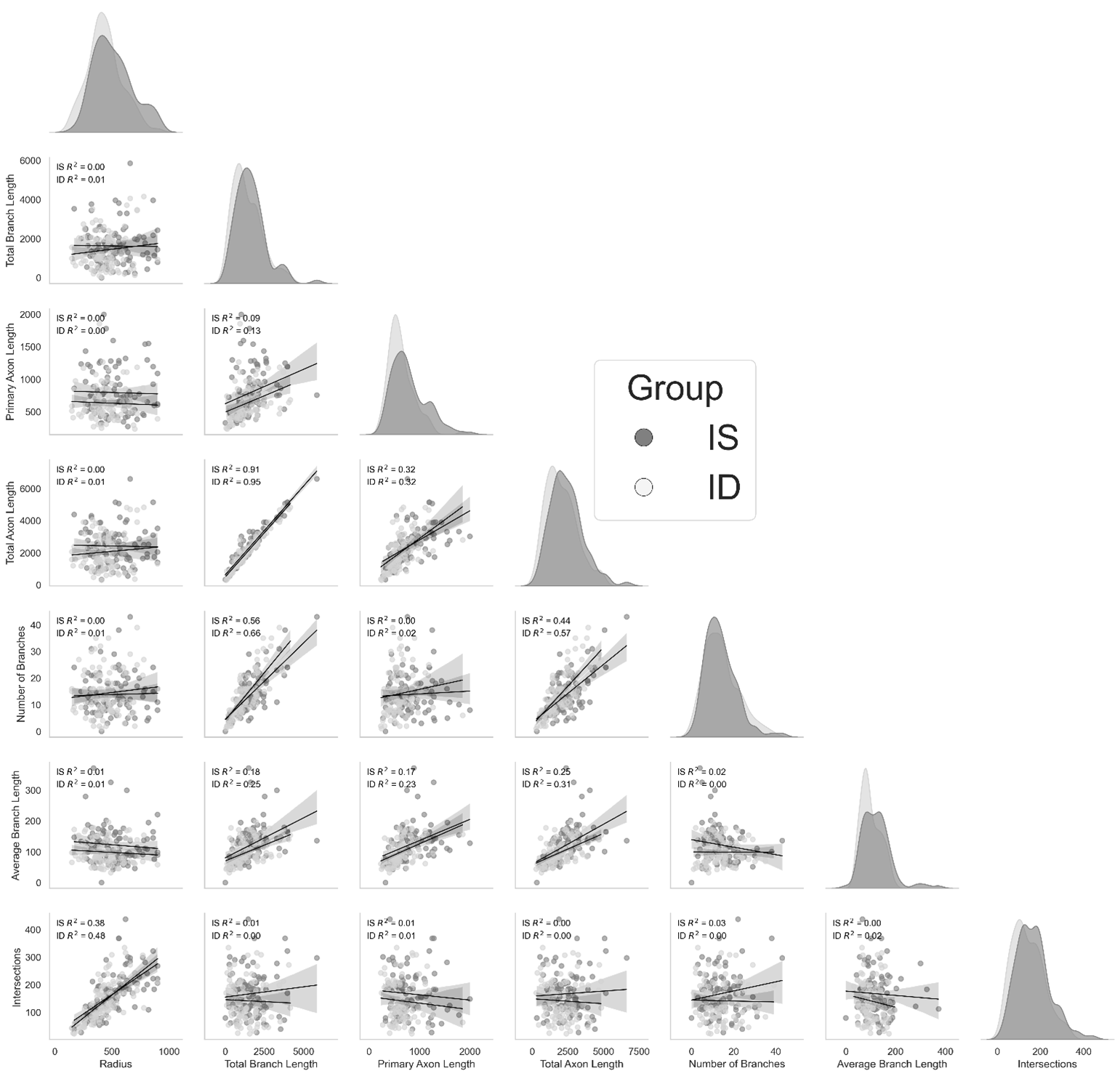

**Supplementary Figure 2: Pairwise regression analysis reveal no differences between IS versus ID neurons.** Heatmap shows matrix of variables along x- and y-axis paired from multivariate regression analysis of the slope differences between IS versus ID. The FDR adjusted p-values are displayed and colored according to the heatmap scale from black (low p-value) to pink (high p-value). Statistical significance was considered when p ≤ 0.05.

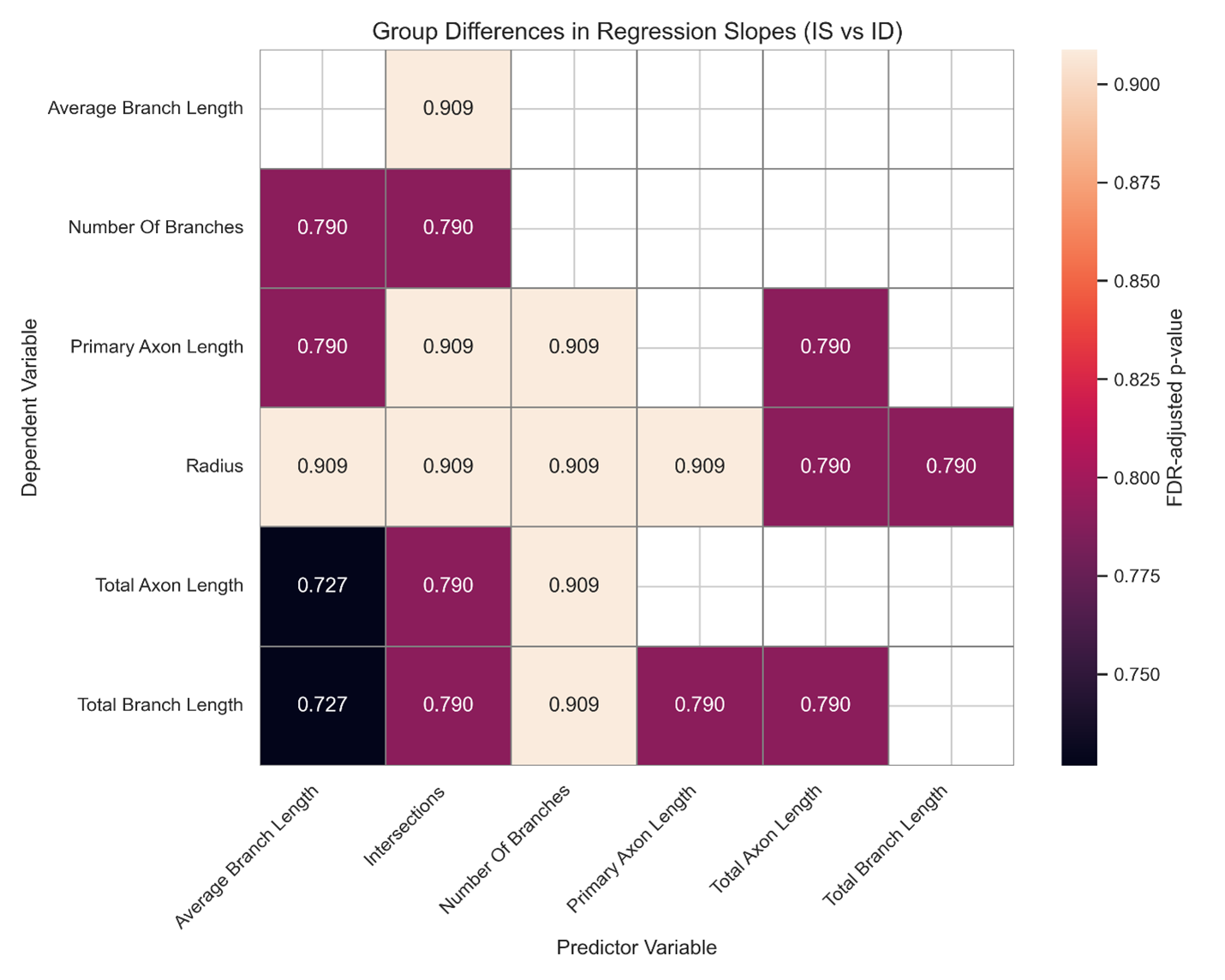

**Supplemental Table 2: Tabulated heatmap statistics show no difference in pairwise regression statistics between IS versus ID neurons.** Table shows output linear regression statistic differences of multivariate pairwise comparisons between IS versus ID**.** Results are organized by row for each variable pair association compared for differences between IS versus ID neurons. The column categories displayed in the table include x- and y- variables, FDR adjusted p-values, effect size, upper and lower confidence intervals and significance (statistical significance considered p≤ 0.05).

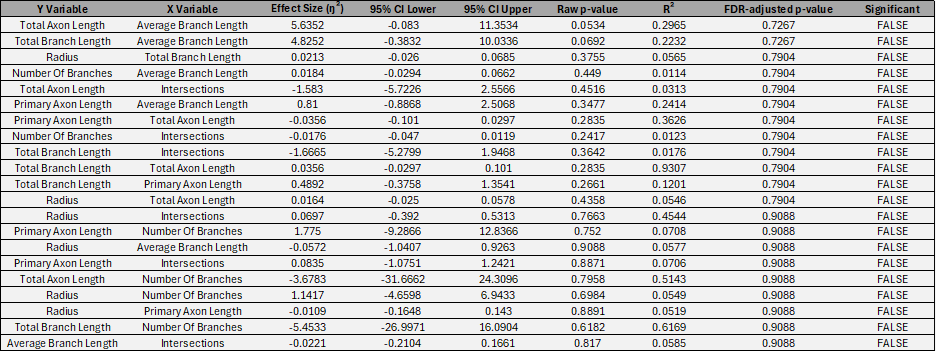

**Supplemental Table 3: Primary axon and average branch lengths are not corrleated with number of branches based on ID status at 7DIV.** Table shows output linear regression statistics of pairwise comparisons between variable pairs in multivariate analysis**.** Statistical results are organized by row for each variable pair association in the IS (dark gray) and ID (light gray) groups. The column categories displayed in the table include x- and y- variables, slope, intercept, R^2^ values, raw p-values, FDR adjusted p-values, and significance (statistical significance considered p≤ 0.05).

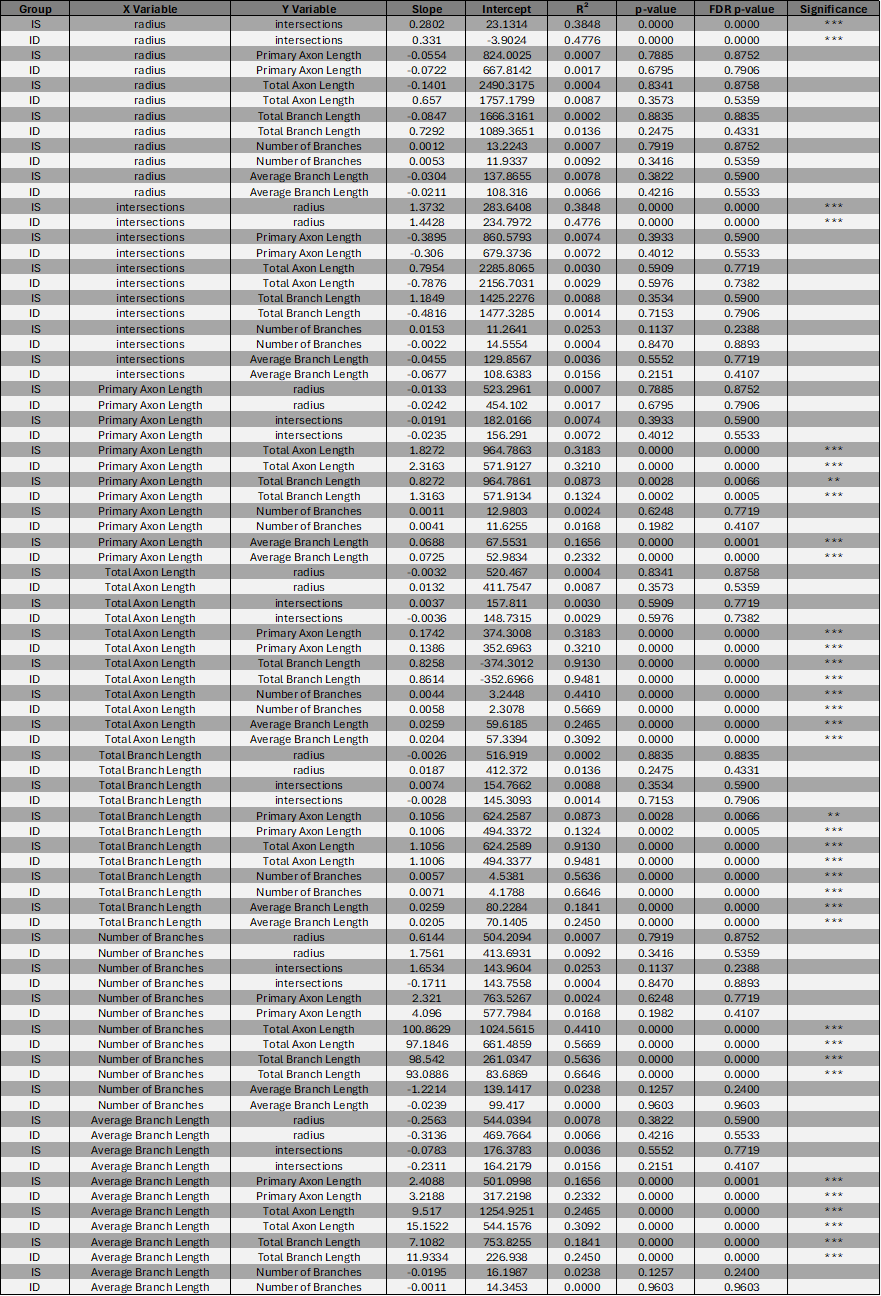
